# Design of complicated all-α protein structures

**DOI:** 10.1101/2021.07.14.449347

**Authors:** Koya Sakuma, Naohiro Kobayashi, Toshihiko Sugiki, Toshio Nagashima, Toshimichi Fujiwara, Kano Suzuki, Naoya Kobayashi, Takeshi Murata, Takahiro Kosugi, Rie Koga, Nobuyasu Koga

## Abstract

A wide range of de novo protein structure designs have been achieved, but the complexity of naturally occurring protein structures is still far beyond these designs. To expand the diversity and complexity of de novo designed protein structures, we sought to develop a method for designing “difficult-to-describe”α-helical protein structures composed of irregularly aligned α-helices like globins. Backbone structure libraries consisting of a myriad of α-helical structures with 5- or 6-helices were generated by combining 18 helix-loop-helix motifs and canonical α-helices, and five distinct topologies were selected for de novo design. The designs were found to be monomeric with high thermal stability in solution and fold into the target topologies with atomic accuracy. This study demonstrated that complicated α-helical proteins are created using typical building blocks. The method we developed would enable us to explore the universe of protein structures for designing novel functional proteins.

---

Many naturally occurring protein structures are complicated, lacking distinguishable symmetry and regularity. Prominent examples of such complicated proteins are globin-fold structures with eight irregularly packed α-helices; Kendrew referred to the tertiary arrangement of the secondary structures as being difficult to describe in simple terms (*1*) (Fig. 1A). In most parts of globin fold structures, two helices adjacent in the sequence are connected crosswise rather than hairpin-like, and the helix-helix packings deviate from the canonical patterns (*2, 3*); this fold does not include internal structural repeats such as α-solenoids (*4, 5*). These asymmetric, irregular, and non-repetitive secondary structure arrangements make it difficult to simply describe globin structures, and so are many naturally occurring proteins.

A wide range of all-α protein structures have been designed, but the designs have been limited to simple and ordered structures consisting of almost parallelly aligned α-helices, such as coiled-coil, bundle, and barrel structures (Fig. 1, B-D, and fig. S1) (*5–27*). Jacobs *et al.* attempted to design α-helical proteins with more variety (*15*), but their designs were still bundle-like (the two designs with five α-helices in Fig. 1B). However, the complexity of tertiary structures underlies various protein functions; proteins with complicated tertiary structures can have diverse and heterogenous molecular surfaces of active sites, thus enabling specific interactions with binding partners or substrates. Therefore, the ability to create protein structures with complicated secondary-structure spatial arrangements like globins would contribute to the custom design of various functional proteins. In this study, we sought to develop a computational method for designing complicated all-α structures.

**Fig. 1.**
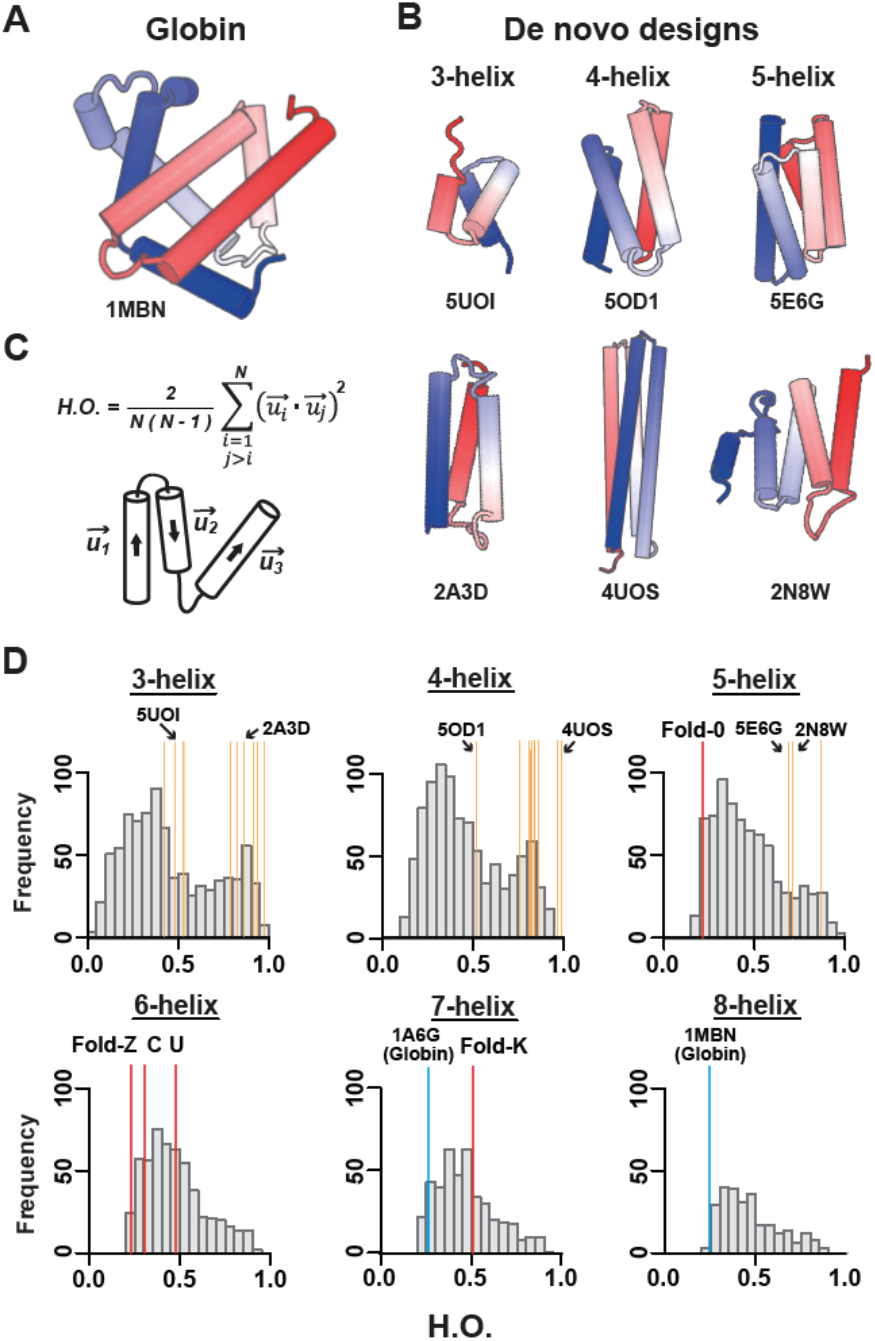
Comparison of the structural complexities of naturally occurring and de novo designed proteins. **(A)** Structures of myoglobin and **(B)** representative de novo designed all-α proteins (the N and C-terminals are colored in blue and red, respectively; and the characters represent PDB IDs). The α-helices in the globin structure are irregularly aligned, whereas those of the de novo designs are almost parallelly aligned. **(C)** The order parameter capturing the complexities of α-helical proteins, Helix Order (H.O.). H.O. is defined by the average of inner products between helix orientation vectors, *u_i_*, for all pairs of *N* α-helices (*62*). Higher values indicate more ordered and lower values, more complicated. **(D)** H.O. distributions for naturally occurring and de novo designed proteins with three to eight α-helices. Whereas naturally occurring all-α proteins show broad distributions irrespective of the number of constituent α-helices, previous de novo designed all-α proteins indicated by yellow-ocher bars show relatively higher values in the distributions (see fig. S1 for details of the previous designs). Notably, globin structures indicated by blue bars have quite low values. The all-α proteins created in this study, indicated by red bars, have lower values than the previous designs.

The major obstacle in designing complicated all-α topologies with irregularly aligned α-helices is attributed to the difficulty in imagining such topologies and drawing their backbone blueprints. This is different from the design of αβ-proteins; the topologies are mainly described by β-strand arrangements, and the backbone blueprints involving lengths of secondary structures and loops were derived from a set of rules relating local backbone structures of a few successive secondary structure elements to the preferred tertiary motifs (*28*). Therefore, we attempted to explore all-α topologies, not by preparing them a priori but by generating backbone structure topologies through the combinatorial enumeration of tertiary building blocks (Fig. 2). Moreover, the tertiary building blocks were selected from those typically observed in nature, so that the generated backbone structures are likely to be designable. The question is whether complicated all-α topologies can be generated from typical building blocks.

**Fig. 2.**
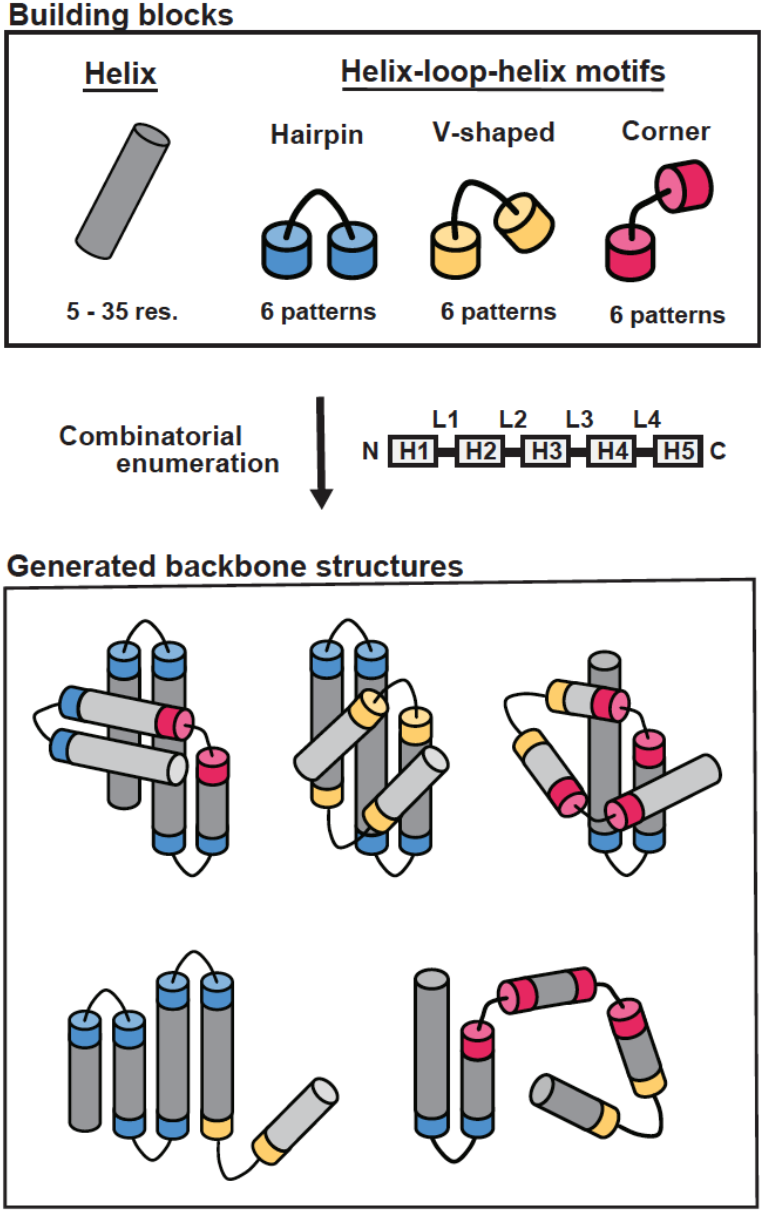
Strategy for building α-helical backbone structure topologies. (Top) Building blocks for generating backbone structures. Canonical α-helices and three types of HLH tertiary motifs typically observed in nature, hairpin (h), v-shaped (v), and corner (c), are used. Helices range from 5 to 35 residues and each motif type comprises 6 patterns (see Fig. 3A). The motif types were classified on the basis of the bending angle between the constituent helices in HLH motifs. (Middle) Secondary-structure element ordering to build α-helical proteins with five helices. According to the ordering, globular backbone structures without steric clashes are exhaustively explored by combining the building blocks, with the constraint of total residue length. (Bottom) Examples for generated α-helical backbone structure topologies. Poorly packed structures (lower) are discarded, whereas globularly folded structures (upper) are collected.

We first attempted to collect a set of helix-loop-helix (HLH) tertiary motifs that are typically observed in nature as building blocks. The HLH units consisting of two α-helices and the connecting loop of one to five residues in length were extracted from naturally occurring proteins, then clustered into 18 subgroups based on the five-dimensional feature vectors representing the HLH tertiary geometries (*29*) (figs. S2 and S3, and supplementary methods). The representative 18 HLH motifs corresponding to each cluster density peak exhibited a broad range of bending angles between two helices, such as left- or right-handed helix-turn-helix, helix-corner-helix, and kinked helices (Fig. 3A and figs. S3 and S4; the amino acid preference for each motif is shown in fig. S5), which can be classified into three classes according to the magnitude of the bending angle: hairpin (h), v-shaped (v), and corner (c). The 18 representative HLH motifs were used as building blocks for generating α-helical backbone structure topologies.

**Fig. 3.**
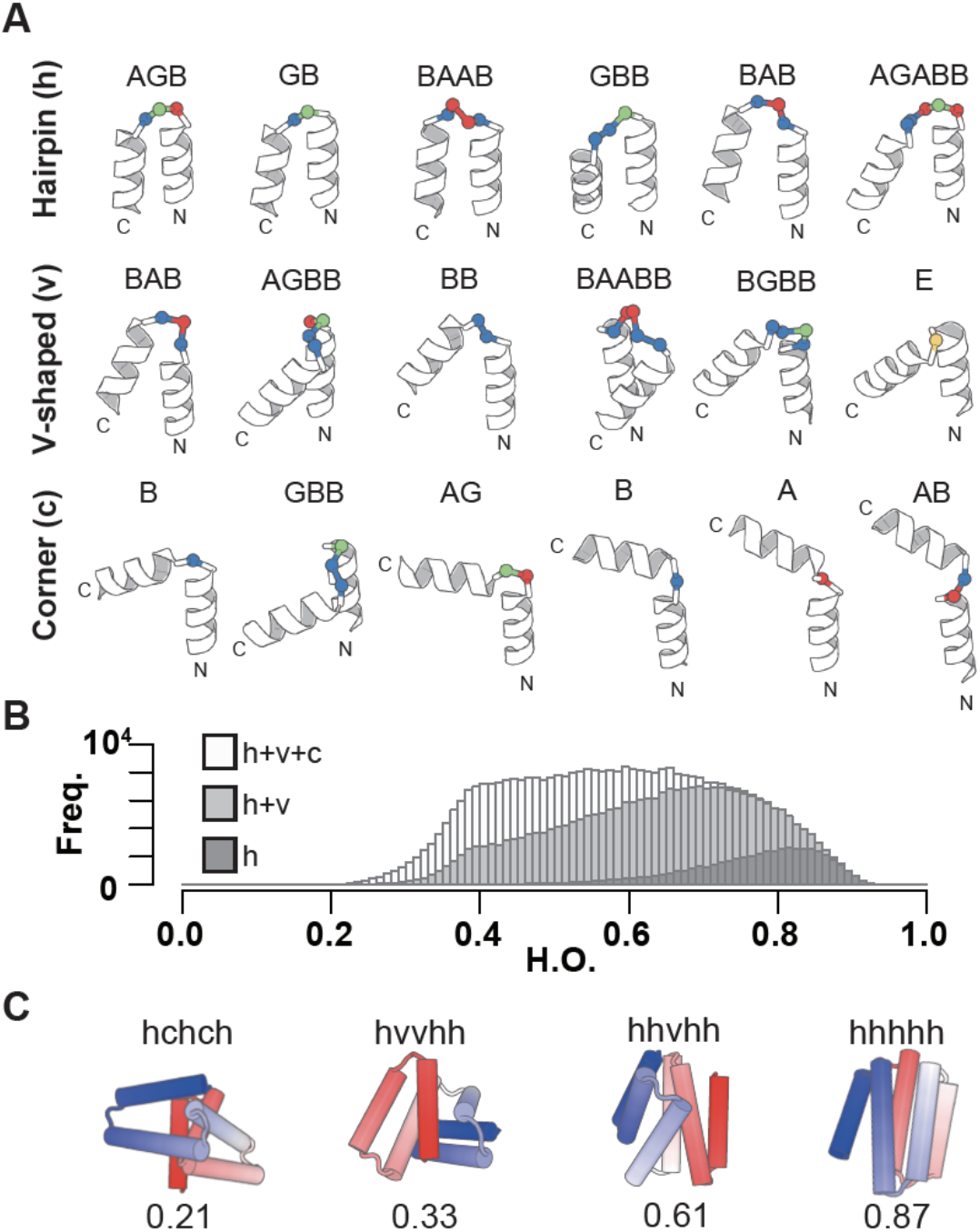
18 HLH tertiary motifs and generated α-helical backbone structures. **(A)** Identified 18 HLH tertiary motifs typically observed in nature. The motifs are classified based on the bending angle between the two helices in the motifs: hairpin (h), v-shaped (v), and corner (c), which are presented in order of the magnitude of the bending angle, with the ABEGO backbone torsion pattern for the connecting loop. The residues with the backbone torsion angle, A, B, E, and G, in the ABEGO torsion representation (“A” corresponds to the right-handed α-helix region in the Ramachandran map, “B” to the β-strand region, “E” to the extended region with a positive phi angle, and “G” to a left-handed α-helix) are shown in red, blue, yellow, and green, respectively. **(B)** H.O. distributions for generated backbone structures with six helices. The black, grey, and white bars respectively represent the distributions for the ensemble generated using only hairpin motifs (h), hairpin and v-shaped (h+v) motifs, and all three motifs (h+v+c). Incorporation of v-shaped and corner loops yields lower H.O. structures. **(C)** Examples for the generated backbone structures. The used motif type strings and the H.O. values are indicated for each structure.

Next, we investigated whether complicated topologies are produced using these typical tertiary motifs. Helical backbone structures composed of 5- and 6-helices were built with 90 and 110 residues in the total length respectively, by combining the set of 18 HLH motifs and canonical α-helices ranging from 5 to 35 residues. The backbone structures were generated by enumerating all the combinations and selecting compact and steric-clash-free structures (see supplementary methods): 1,159,937,910 five-helix and 20,878,882,380 six-helix structures were generated, and 1,899,355 and 380,869 structures were then selected for each. The resulting topologies exhibited a broad spectrum ranging from helical bundle-like to complicated globular structures, demonstrating that complicated α-helical topologies are created from the typical tertiary motifs and canonical α-helices (white bar in Fig. 3B, Fig. 3C, and fig. S6). Moreover, we found that the complexities of the generated topologies increase, as tertiary motifs with larger bending angles are included (black, gray, and white bars in Fig. 3B). These results highlight the importance of corner-type motifs (*30*) in building complicated α-helical topologies.

From the generated myriad backbone structure topologies, we manually selected five for de novo design, H5_fold-0, H6_fold-C, H6_fold-Z, H6_fold-U, H7_fold-K (the Arabic numeral after ‘H’ indicates the number of helices) (Fig. 4 and fig. S7). We first selected three topologies exhibiting extremely low helix order (H.O.) values (See Fig. 1C for the definition): H5_fold-0, H6_fold-C, and H6_fold-Z (Fig. 1D). Next, to test whether all identified HLH motifs could be used for de novo design, we selected H6_fold-U and H7_fold-K, which include all of the HLH motifs not used in the first three and still exhibit lower H.O. values (Fig. 1D). For all target folds except H5_fold-0, the lengths of the terminal helices were manually elongated to ensure sufficient packing interactions. None of these backbone structures is similar to any known protein structures; H5_fold-0, H6_fold-C, H6_fold-Z, and H6-fold-U show a TM-score < 0.6, using TM-align (*31*) against the ECOD database (*32*), and H6_fold-K shows a score of 0.610, with a structure of e2bnlA1 (fig. S8). The details of the selected topologies are described in the supplementary text. For each backbone structure, amino acid sequences were designed through iterations of fixed-backbone sequence optimization and fixed-sequence structure optimization using Rosetta design calculations (*33, 34*). Designs with low energy, tight core packing, and high compatibility between local sequences and structures (*28*) were selected, and their energy landscapes were explored by 10,000 independent Rosetta *ab initio* structure prediction simulations starting from an extended conformation (*35*). The designs showing funnel-shaped energy landscapes (Fig. 5A) were selected for the experimental characterization.

**Fig. 4.**
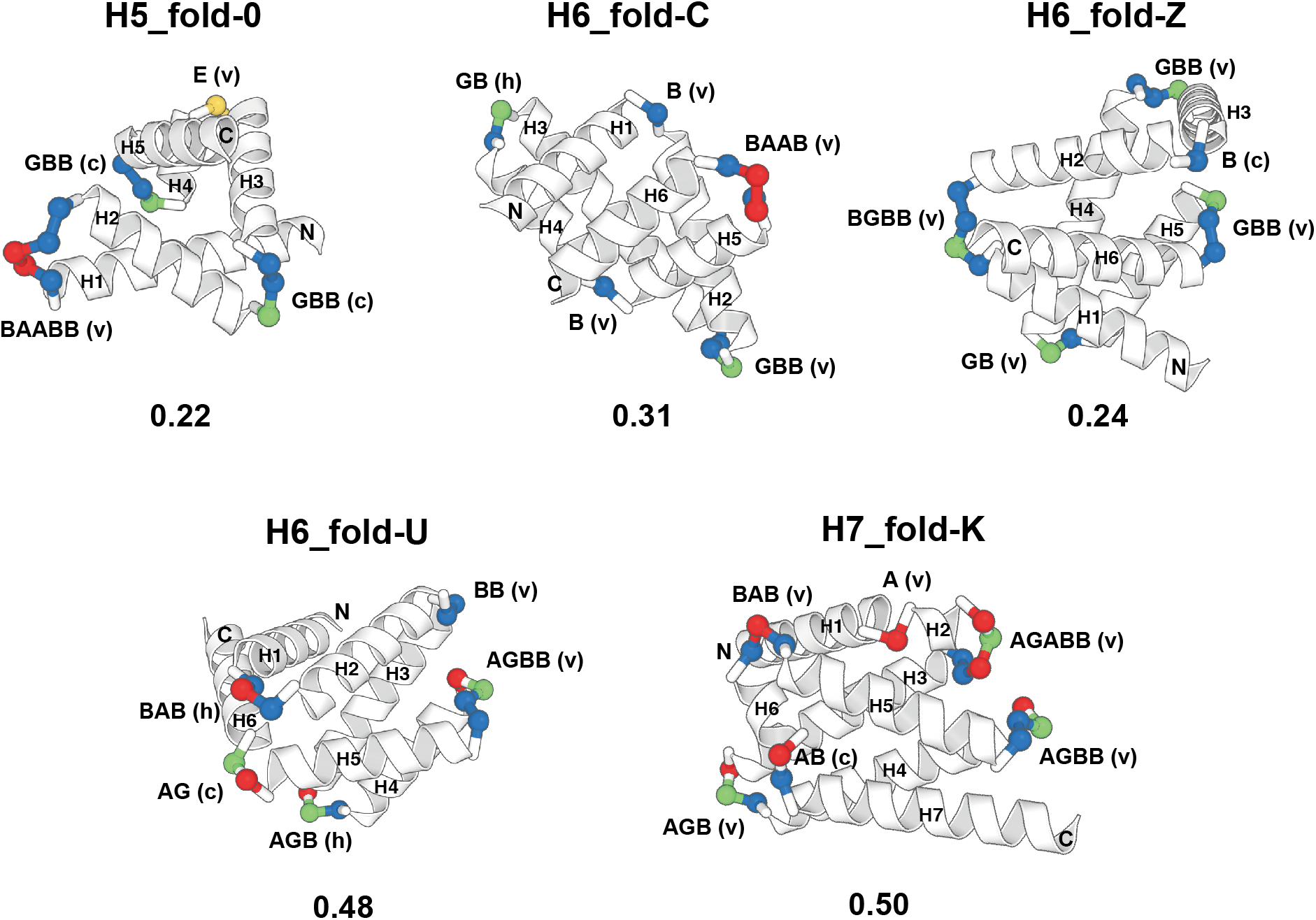
Backbone structures for the five design target topologies. Design target backbone structures. H1-7 represents the first to seventh helices. The letter string next to a loop indicates the ABEGO torsion pattern and the character within a bracket indicates the motif type. The loop residues are colored in the ABEGO torsion representation, same as Fig. 3A. The H.O. values are indicated for each structure.

We obtained synthetic genes encoding 10 designs for H5_fold-0, 7 for H6_fold-C, 7 for H6_fold-Z, 8 for H6_fold-U, and 8 for H6_fold-K. Some designs (H6_fold-Z: 2, H6_fold-U: 1, H7_fold-K: 2) have weak sequence similarity to known proteins with blast *E*-value < 0.005, but the structures are unknown (table S11). The proteins were expressed in *E. coli* and purified using a Ni-NTA affinity column. The purified proteins were then characterized by circular dichroism (CD) spectroscopy and size-exclusion chromatography combined with multi-angle light scattering (SEC-MALS). For all target design topologies, 34 of 40 designed proteins were found to be well-expressed and highly soluble, and showed CD spectra typical of α-helical proteins; 27 out of the 34 designs were found to be monomeric by SEC-MALS (tables S2-6). Furthermore, the monomeric designs were characterized by ^1^H-^15^N heteronuclear single quantum coherence (HSQC) NMR spectroscopy, and 22 designs showed well-dispersed sharp peaks (tables S2-6). The experimental results for all the designs are summarized in table S1. For each topology, we selected one monomeric design with well-dispersed sharp NMR peaks for NMR structure determination (Fig. 5 and fig. S9). All the designs were found to be highly stable from thermal denaturation up to 170 °C by CD (Fig. 5b, c). The NMR structures were solved at high quality using MagRO-NMRViewJ (*36, 37*) (supplementary text, fig. S10 and table S7), and the solved structures were consistent with the design models (Fig. 6 and table S9). For H5_fold-0, one of the designs was solved by X-ray crystallography and was nearly identical to the design model except for the domain swapping in the crystallized condition (Fig. 6 and table S8). Despite the inclusion of non-canonical helix-helix packing arrangements in each design, the side-chains from distant α-helices were found to coherently be packed to constitute a single hydrophobic core similar to the design model. Notably, the bulky hydrophobic side-chains from the loops and neighboring α-helices also contributed largely to the core: they spiked the core and pinned the loops to the target conformations (fig. S11). We compared the loop geometries of all HLH motifs at the ABEGO level in the design models and experimental structures (fig. S12 and table S10). All but one loop geometries of the experimental structures agreed with those of the design models. These results indicate that the difficult-to-describe α-helical proteins are designable with typical building blocks.

**Fig. 5.**
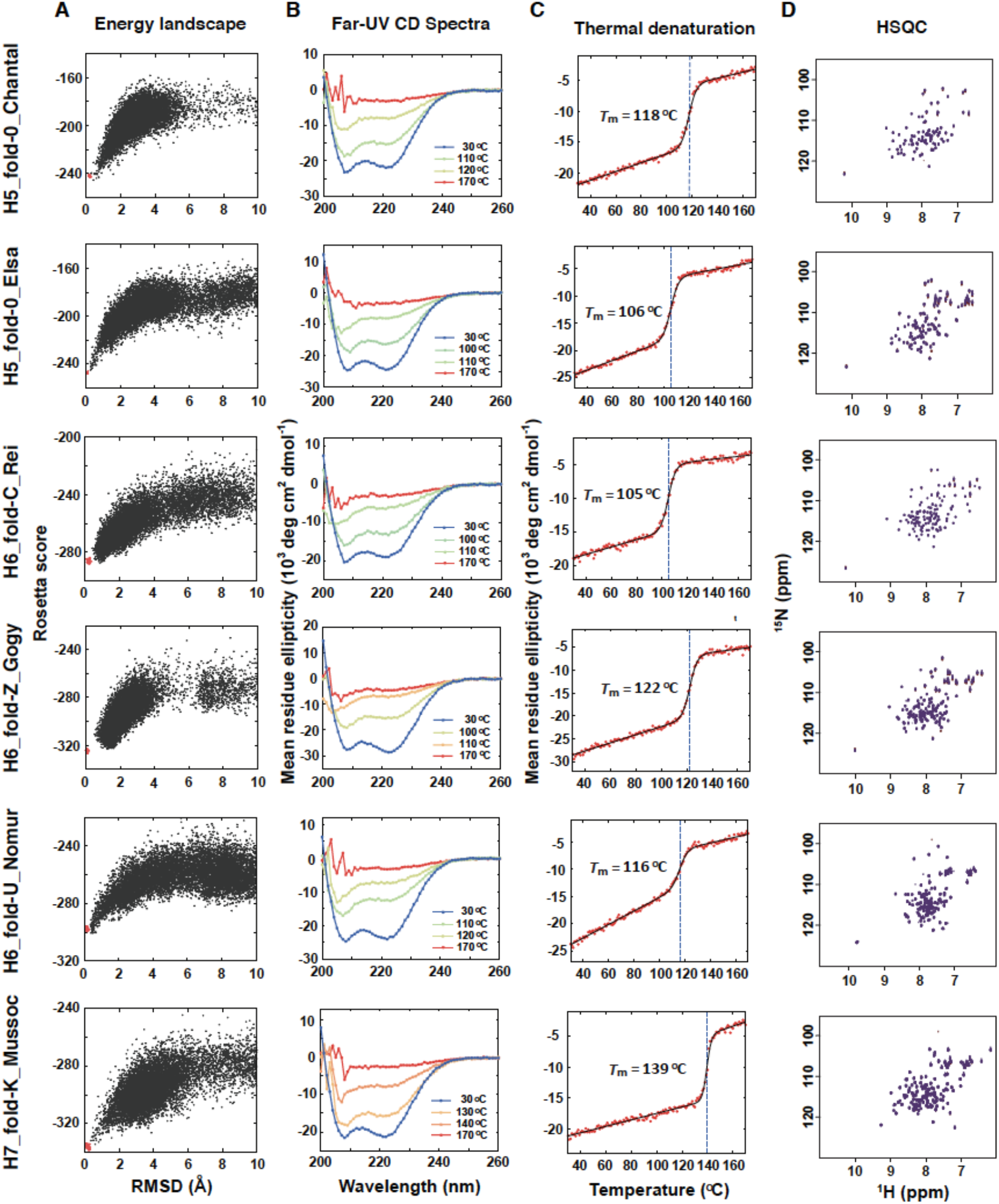
Characterization of designed proteins. **(A)** Energy landscapes from Rosetta *ab initio* structure prediction simulations. The *y*-axis represents Rosetta all-atom energy and the *x*-axis represents the Cα root mean square deviation (RMSD) from the design model. Black points represent the lowest energy structures obtained in independent Monte Carlo structure prediction trajectories starting from an extended chain for each sequence; red points represent the lowest energy structures obtained in trajectories starting from the design model. **(B)** Far-ultraviolet circular dichroism (CD) spectra at 30 C, the temperatures close to the melting temperature *T_m_*, and 170 C. **(C)** Thermal denaturation measured at 222 nm. The data were fitted to a two-state model (solid line) to obtain the *T_m_*. **(D)** Two-dimensional ^1^H-^15^N HSQC spectra at 25 C and 600 MHz.

**Fig. 6.**
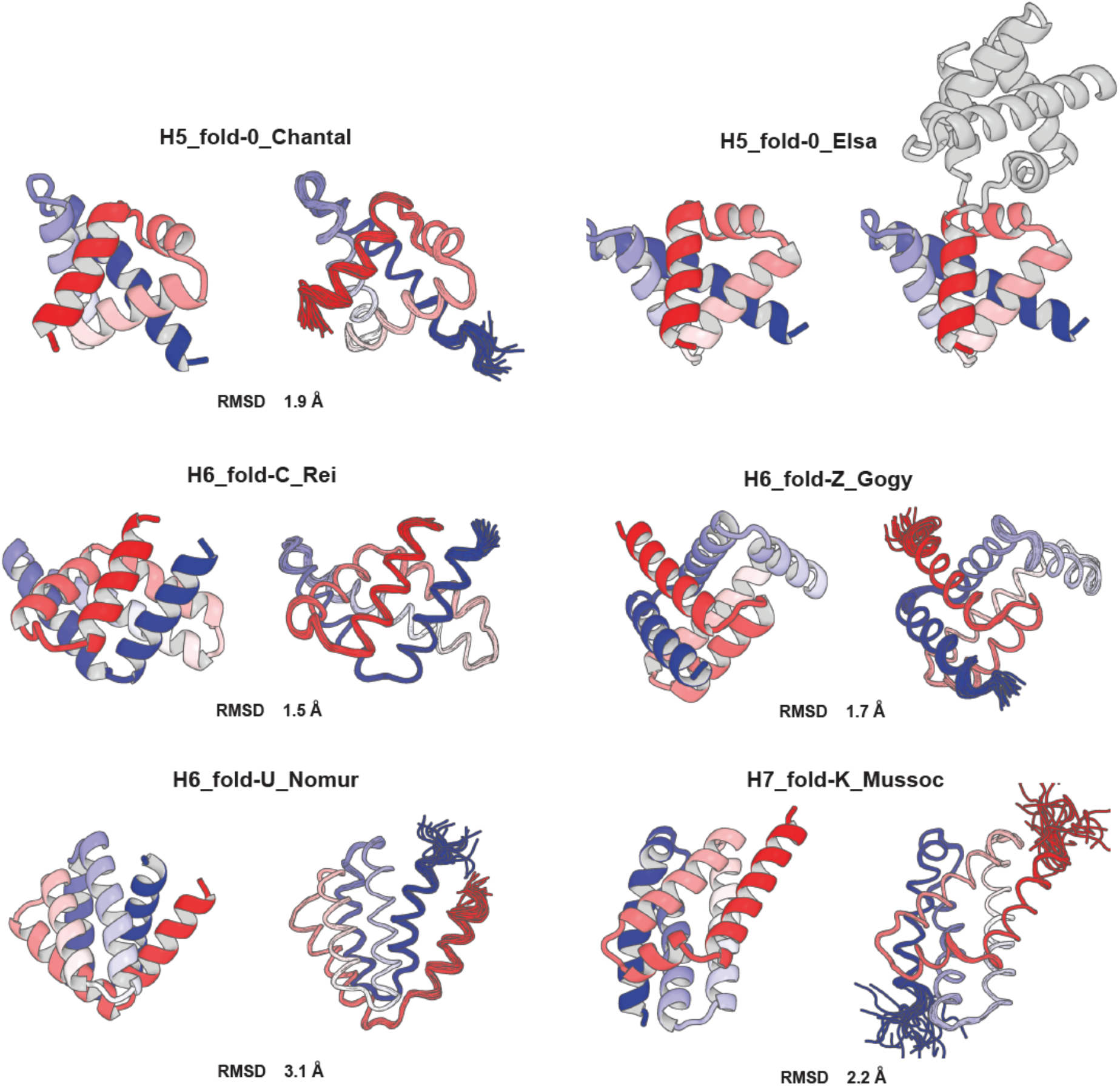
Comparison of computational models with experimentally determined structures. Design structures (left) and NMR structures (a crystal structure for H5_fold-0_Elsa) (right); the N and C-terminals are colored in blue and red, respectively. The Ca root mean square deviation (RMSD) between them is indicated.

*De novo* designs of α-helical proteins have focused on structures consisting of parallelly aligned α-helices (Fig. 1), many of which are based on helical structure models such as the helical wheel (*38*) and Crick’s parameterization (*39*). We sought to develop a computational method for designing difficult-to-describe α-helical protein structures. We demonstrated that a wide range of globular all-α backbone structure topologies from bundle-like to complicated are generated by combining 18 identified typical HLH motifs and canonical α-helices. This approach is regarded as the reverse of blueprint-based design: design target topologies are searched by the combinations of HLH motifs in this approach, whereas design target topologies are predetermined and then local backbone structures favoring the topologies are selected in blueprint-based design. The key to building complicated α-helical topologies is to include HLH motifs with larger bending angles such as corner-type motifs. We succeeded in designing complicated α-helical protein structures with five distinct topologies, three of which exhibited structural complexities comparable to the globin fold. The design success rate was as high as that of previous de novo designs, and the design exhibited high solubility and thermal stability, similarly to previous designs (*28, 40–44*). Moreover, the loop geometries of almost all HLH motifs were formed as designed, which must have enabled the designed proteins to fold into the target topologies. These results suggest that a variety of α-helical protein structures generated using our strategy are designable. Our design method enables to explore the helical protein universe beyond bundle-like proteins (fig. S6).

The complicated α-helical proteins provide diverse and heterogeneous molecular surfaces for engineering functions such as binding, enzymatic activity, and self-assembly into symmetric oligomers. Most naturally occurring proteins generate diverse molecular surfaces by changing their loop conformations. This can be attributed to the protein evolution strategy of gene insertion or deletion, which enables to locally change the conformations, but it is likely difficult to globally rearrange the spatial arrangements of secondary structures. The computationally generated myriad of diverse proteins, presumably with high solubility and stability, coupled with the recently developed massive gene synthesis (*45, 46*) and parallel high-throughput screening (*17, 18, 26, 47*), should make it possible to create proteins with optimal structures for specific functions (*17, 26*).

## Methods

### Analysis of all-α protein structures for de novo designed and naturally occurring proteins

22 de novo designed all-α protein structures were collected from Protein Data Bank (PDB). To this end, de novo designs were searched by the keyword ‘de novo’ or ‘de-novo’ in PDB as of November, 2020, and then all-α structures containing no β-strands were extracted based on the secondary structure assignments by the DSSP algorithm (*48*) (for the PDB structures including multiple chains or NMR models, the first chain or model was used). The following four classes of de novo designed proteins were excluded from the dataset. (1) Designed proteins created based on backbone structures of naturally occurring proteins, and those with sequence similarity higher than 0.90 (as an exception, the three-helix bundle structure designs (PDB code: 6DS9 and 2A3D) were both included because of their structural dissimilarity). (2) Assemblies composed of one or two α-helices (e.g.,3R3K and 1U7J). (3) Repetitive structures like α-solenoids (e.g., 1MJ0 and 5K7V). (4) Membrane proteins.

For naturally occurring all-α protein structures, 7352 representative structures found in the mainly-α class in the CATH database (*49*) with sequence identity less than 40% were used. For calculating the H.O. values of the collected structures, secondary structure elements and loops were assigned by DSSP (*48*) (α-helices are defined for the residue regions composed of at least five successive residues assigned as ‘H’ by the DSSP calculation). Note that the secondary structure assignments by DSSP are not always consistent with those originally defined by the authors. For example, the number of α-helices in the structures (PDB codes: 1P68 and 4TQL) respectively designed with three and four α-helices were defined as four and five due to partially distorted α-helices.

### Clustering of HLH units using the five features representing a HLH geometry

13,667 HLH structures were extracted from 7,280 X-ray structures (secondary structures were assigned by DSSP (*48*)), obtained from the PISCES server (*50*), with resolution ≤ 2.5Å, R-factor ≤ 0.3, sequence length more than or equal to 40, and ≤ 25% sequence identity. We then classified the HLH structures by their loop residue lengths and extracted 13,510 HLH structures in total with loop of one to five residues in length. The extracted HLH structures were clustered for each loop length from one to five using the density clustering algorithm (*29*) (fig. S3), with the five features representing a HLH geometry (fig. S2).

### Building backbone structures

α-helical backbone structures were built using Rosetta by exhaustive sampling for the conformations with steric-clash free (Rosetta vdw score < 4.0 using the weight value, 0.1) and smaller radius of gyration (< 14 Å) by combining canonical α-helices ranging from 5 to 35 residues (backbone torsion angles, phi, psi, and omega, were set to −60.0, −45.0, 180.0 respectively) and the identified 18 helix-loop-helix (HLH) motifs (see main text and Fig. 3A), with length constraints of 90 and 110 residues for the five- and six-helix proteins, respectively. For generating five-helix structures, 64,440,995 steric-clash free four-helix structures with 70 residues were first generated, and then an α-helix with 18 types of connecting loops was appended to the C-terminal of the generated four-helix structures so that the total length becomes 90 residues. For generating six-helix structures, an α-helix with 18 types of connecting loops was appended to the N-terminal of the generated five-helix structures so that the total length becomes 110 residues. From these structures, the globular five- and six-helix structures were collected based on the radius of gyration.

### Expression and purification of designed proteins

The genes encoding the designed sequences were synthesized and inserted into pET21b vectors. The whole plasmid constructs were purchased from FASMAC or Eurofins Genomics. The target proteins were overexpressed by IPTG induction in *E. coli* BL21 Star (DE3) cells cultured in MJ9 minimal media including ^15^N ammonium sulphate as the sole nitrogen source and ^12^C glucose as the sole carbon source (*51*). The expressed uniformly (*U*-)^15^N-labeled proteins with a 6xHis tag at the C-terminus were purified by Ni-affinity columns. The purified proteins were then dialyzed against PBS buffer, 137 mM NaCl, 2.7 mM KCl, 10 mM Na_2_HPO_4_, 1.8 mM KH_2_PO_4_, at pH 7.4; this buffer was used for all the experiments except NMR structure determination. The expression level, solubility, and purity of each designed protein were evaluated by SDS-PAGE. To further confirm them, the samples were analyzed by mass spectroscopy (Bruker Daltonics REFLEX III and Thermo Scientific Orbitrap Elite).

### Experiments to identify designed proteins exhibiting folding ability

The following three experiments were conducted to evaluate the folding ability of designed sequences: circular dichroism (CD) spectroscopy, size exclusion chromatography with multi-angle light scattering (SEC-MALS), and ^1^H-^15^N hetero-nuclear single quantum coherence (HSQC) nuclear magnetic resonance (NMR) spectroscopy. Tables S2-6 show the results of the evaluations for each designed sequence for each fold.

### CD spectroscopy under 1-bar pressure

Far-UV CD spectra was measured to study whether the designs show the characteristic spectra of α-helical proteins, by scanning from 260 to 200 nm at 20 °C for ~15 μM protein samples in PBS buffer on a JASCO J-1500 KS CD spectrometer. The measurements were performed 4 times and then averaged.

### SEC-MALS

Oligomeric states for the designs in solution were studied by SEC-MALS with miniDAWN TREOS static light scattering detector (Wyatt Technology Corp.) combined with a HPLC system (1260 Infinity LC, Agilent Technologies) with a Superdex 75 increase 10/300 GL column (GE Healthcare). After the equilibration of the column with PBS buffer, 100 μL of the samples after purification by Ni-affinity columns were injected. The absorbance at 280 nm was measured by the HPLC system to give the protein concentrations and intensity of light scattering at 659 nm was measured at angles of 43.6°, 90.0°, 136.4°. These data were analyzed by the ASTRA software (version 6.1.2, Wyatt Technology Corp.) using a change in the refractive index with concentration, a *dn/dc* value, 0.185 ml/g, to estimate the molecular weight of dominant peaks.

### ^1^H-^15^N HSQC NMR spectroscopy

Whether the designs fold into well-packed structures or not was evaluated by ^1^H-^15^N HSQC 2D-NMR spectroscopy. The purified protein samples were concentrated to 0.2-1.0 mM, and mixed with their 10% volume of D_2_O. The measurements were performed at 25°C on a JEOL JNM-ECA 600 MHz spectrometer, and data were analyzed by JEOL Delta (version 5.3.1).

### High-pressure CD spectroscopy for melting temperature (*T_m_*) estimation

For the designs that were evaluated to have the folding ability in the above experiments (one design for each target topology was selected), thermal denaturation was studied by using high-pressure CD spectroscopy. JASCO J-1500 KS CD spectrometer was equipped with additional pressure instruments so that temperature of the solution samples can be scanned from 30 to 170 °C under 10 bar. Temperature was increased 1 °C per minute for ~15 μM protein samples. Fixed wavelength measurements at 222 nm were performed at every 1 °C, and wavelength scanning measurements (260 to 200 nm) were performed at 30, 40, 60, 80, 90, 100, 110, 120, 130, 140, 150, 160, and 170 °C. *T_m_* was estimated by non-linear fitting to thermal denaturation CD curve at 222 nm. The non-linear least-squares analysis was performed by *nls* function in R language, given a two-state unfolding and linear extrapolation model. After this fitting, we obtained *T_m_* at which the estimated populations of folded and unfolded states get equal.

### Sample preparation for NMR structure determination

The most promising design for each target topology was overexpressed by IPTG induction in *E. coli* BL21 Star (DE3) cells cultured in MJ9 minimal media containing ^15^N ammonium sulphate as the sole nitrogen source and ^13^C glucose as the sole carbon source (*51*). The expressed *U*-^15^N,*U*-^13^C-enriched proteins were purified by Ni-affinity columns, and dialyzed against PBS buffer. The protein samples were further purified by gel filtration chromatography on an ÄKTA Pure 25 FPLC (GE Healthcare) using a Superdex75 or Superdex75 increase 10/300 GL column (GE Healthcare), which also replaces the PBS buffer at pH 7.4 with the customized buffer for NMR spectroscopy. The following 95% H_2_O/5% D_2_O buffer conditions for each sample were used: 100 mM NaCl, 5.6 mM Na_2_HPO_4_, 1.1 mM KH_2_PO_4_, at pH 7.4 for H5_fold-0_Chantal; 50 mM NaCl, 5.5 mM Na_2_HPO_4_, 4.5 mM KH_2_PO_4_, at pH 6.9 for H6_fold-C_Rei; 50 mM NaCl, 3.2 mM Na_2_HPO_4_, 4.5 mM KH_2_PO_4_, at pH 6.5 for H6_fold-Z_Gogy; 155 mM NaCl, 3.0 mM Na_2_HPO_4_, 1.1 mM KH_2_PO_4_, 10 μM EDTA, 0.02% NaN_3_, cOmplete protease inhibitor cocktail (Roche), at pH 7.4 for H6_fold-U_Nomur; and 155 mM NaCl, 3.0 mM Na_2_HPO_4_, 1.1 mM KH_2_PO_4_, at pH 7.4 for H7_fold-K_Mussoc.

### Solution structure determination by NMR

#### NMR measurements

NMR measurements were performed on Bruker AVANCE III NMR spectrometers equipped with QCI cryo-Probe at 303K. The spectrometers with 600, 700, and 800 MHz magnets were used for the signal assignments and NOE related measurements, while 700, 900, and 950 MHz ones, for residual dipolar coupling (RDC) experiments. For the signal assignments, 2D ^1^H-^15^N HSQC (echo/anti-echo), ^1^H-^13^C Constant-Time HSQC for aliphatic and aromatic signals, 3D HNCO, HN(CO)CACB, and 3D HNCACB for backbone signal assignments, while BEST pulse sequence was applied to the triple resonance measurements for H6_fold-C_Rei. For structure determination, 3D ^15^N-edited NOESY, 3D ^13^C-edited NOESY for aliphatic and aromatic signals (mixing time = 100 ms) were performed. For H6_fold-U_Nomur, additional 3D HN(CA)CO, HN(CO)CA, HNCA, HBHA(CO)NH, HBHANH, H(CCCO)NH, CC(CO)NH, 3D ^13^C-HSQC (^13^C-t1) NOESY ^13^C-HSQC, 3D ^13^C-HSQC (^13^C-t1) NOESY ^15^N-HSQC and 4D ^13^C-HSQC NOESY ^13^C-HSQC were measured. Except for 3D-edited NOESY, all the other spectra were performed using Non-Uniform Sampling (NUS) for H6_fold-U_Nomur and H7_fold-K_Mussoc. For NUS, sampling ratio was set at 25% for 3D and 6% for 4D with a fixed ramdom seed. The NUS spectra were reconstructed by IRLS for 3D while IST for 4D spectra with virtual-echo technique (VE) using qMDD tool.

For the RDC experiments, 2D IPAP ^1^H-^15^N HSQC spectra using water-gate pulses for water suppression were measured with or without 6-10 mg/ml of Pf1 phage (ASLA biotec Ltd.). For confirming the positions of ^1^H-^15^N signals in the 2D IPAP ^1^H-^15^N HSQC, 3D HNCO at the identical buffer condition containing Pf1 phage were measured. The α- and β-states of ^15^N signals split by ^1^H-^15^N J-coupling were separately identified for the protein in the isotropic and weakly aligned states, in order to obtain 1-bond residual dipolar coupling (RDC) ^1^*D*_^1^H/^15^N_ values. For the sample H6_fold-U_Nomur, 3D J-HNCO (without ^1^H decoupling for ^15^N evolution) was measured at 25% NUS, which were used for confirming α- and β-states of ^15^N signal positions overlapped in 2D IPAP spectra. 3D J-HN(CO)CA spectrum was also measured for H6_fold-U_Nomur to obtain ^1^*D*_^1^Ha/^13^ca_ for appending an additional number of alignment data at the identical magnetic field and alignment tensor.

#### NMR signal assignments

All NMR signals were identified in a fully automated manner using MagRO-NMRViewJ (upgraded version of Kujira (*36*)), in which noise peaks were filtered by deep-learning methods using Fit_Robot (*37*). FLYA module was used for fully automated signal assignments and structure calculation (*52*) to obtain roughly assigned chemical shifts (Acs), and then trustful ones were selected into the MagRO Acs table. After confirmation and correction of the Acs by visual inspection using MagRO, TALOS+ (*53*) calculations were performed to predict phi/psi dihedral angles, which were then converted to angle constraints for the CYANA format.

#### Structure calculation

Several CYANA (*54*) calculations were performed using the Acs table, NOE peak table and dihedral angle constraints. The Acs table was exported by the MagRO CYANA module, and then the aliased chemical shifts were automatically calculated depending on the spectrum width of responsible NOESY spectra. For dihedral angle constraints, phi and psi, with deviation are derived from TALOS+ prediction using chemical shifts of ^15^N, ^13^C’, ^13^Ca, and ^13^Cβ, with high prediction score noted by “Good”. The minimal angle deviation was set at 20 deg. After several times CYANA calculations, dihedral angle constraints derived from TALOS+ (*53*) revealing large violation for nearly all models in structure ensemble were eliminated.

After the averaged target function of the ensemble reached to less than 2.0 Å^2^, refinement calculations by Amber12 were carried out for 20 models with lowest target functions. The coordinates of final.pdb calculated by CYANA, distance constraints (final.upl), dihedral angle constraints derived from TALOS+ prediction were converted into Amber format and topology file using Sander Tools. Firstly, 500 steps of minimization (250 steps of steepest decent, 250 steps of conjugate gradient) were carried out without electrostatic potential and NMR constraints. Second, MD simulations with the ff99SB force field using implicit water system (0.1 M of ionic strength, 18.0 Å of cut-off) were performed, in which the temperature was gradually increased from 0.0 K to 300.0 K by 1500 steps, followed by the simulation with 28,500 steps at 300.0 K (1.0 fsec time step, Total 30 psec). Finally, 2000 steps for minimization (1,000 steps for steepest decent and 1,000 steps for conjugate gradient) with constraints of distance and dihedral angle were applied at the same condition used in the MD simulations.

#### NMR structure validation

The RMSD values were calculated for the 20 structures overlaid to the mean coordinates for the ordered regions, automatically identified by Fit_Robot using multi-dimensional non-linear scaling (*55*).

The RDC back-calculation was performed by PALES (*56*) using experimentally determined values of RDC. The averaged correlation between the simulated and experimental values was obtained using the signals except the residues on overlapped regions in ^1^H-^15^N HSQC and the ones in low order-parameters less than 0.8 predicted by TALOS+. For the validation of H6_fold-U_Nomur, a lot of signals are overlapped in 2D IPAP-HSQC spectra. To overcome this problem, ^1^J_HN-15N_ split 3D HNCO (without ^1^H-decoupling scheme in ^15^N evolution period) spectra in isotropic and anisotropic states were measured by NUS (25% data point reduction) to obtain signal positions of α- and β-states of ^15^N spins at resolution of 0.3 Hz. ^1^J_Ha-13Ca_ split 3D HN(CO)CA spectra at the same conditions were also measured to obtain ^1^*D*_^1^Ha/^13^Ca_ at resolution of 0.2 Hz. Initially the RDC reproducibility of H6_fold-U_Nomur were examined using separately ^1^D_HN-15N_ and ^1^D_Ha-13Ca_ tables by PALES for all models to confirm that the averaged RMS values are greater than 0.9, and then final RMS values were calculated with two merged tables.

### X-ray structure determination of H5_fold-0_Elsa

#### Sample preparation for X-ray structure determination

The gene encoding the designed sequence of H5_fold-0_Elsa in pET21b vector was digested at the NdeI and XhoI restriction sites and cloned into pET15b-TEV vector with cleavable sites by TEV protease instead of thrombin (original) between the designed sequence and the N-terminal 6xHis tag. Designed protein was expressed in *E. coli* BL21 Star (DE3) cells, and purified by a Ni-affinity column. The N-terminal His tag was then cleaved by TEV protease, and removed through a Ni-affinity column. The protein samples without a His tag were purified by an anion-exchange chromatography (HiTrapQ HP 1 mL column, GE healthcare) followed by gel filtration chromatography (Superdex 75 10/300 GL column) on an ÄKTA Pure 25 FPLC. Mass spectroscopy was performed to confirm that a His tag was successfully cleaved.

To assess the effect of the tag cleavage on the oligomeric state and stability, we performed SEC-MALS and thermal denaturation CD experiments under high pressure for the original and tag-cleaved samples of H5_fold-0_Elsa. The solvent was exchanged to PBS at pH 7.4 before these experiments. The results showed that the tag-cleaved protein was also monomeric and had nearly identical denaturation temperature (the second row in Fig. 5C, 106 °C) as the original sample with the C-terminal His tag (fig. S13, 105 °C), which indicates that the removal of tag and slight differences in flanking amino-acid sequences do not largely change the stability and oligomeric state of the designed protein in solution.

#### Crystallization and X-ray structure determination

The protein samples of H5_fold-0_Elsa at the concentration of 12 mg/mL (1.07 mM) was crystallized in the solution of 0.4 M MgCl_2_, 0.1 M Tris-HCl (pH 7.5), 30% PEG 3350, using the sitting-drop vapor diffusion method at 296K. The obtained crystals were soaked in the solution of 0.4 M MgCl_2_, 0.1 M Tris-HCl (pH 7.5), 30% PEG 3350, and 10% Glycerol, mounted on cryoloops (Hampton Research), flash-cooled, and stored in liquid nitrogen.

X-ray diffraction data of the crystal were collected with BL-1A beamline at Photon Factory (Tsukuba, Japan), and processed to 2.3 Å by XDS (*57*). After phase determination by molecular replacement using the design model by Molrep (*58*) in the CCP4 suite, the molecular model was constructed and refined using Coot (*59*) and Phenix Refine (*60*). TLS (Translation/Libration/Screw) refinement was performed in late stages of refinement. The refined structures were validated with RAMPAGE (*61*). The crystallographic data collection is summarized in table S8.

## Acknowledgments

We thank RIKEN Yokohama NMR Facility for NMR spectra measurements, the Functional Genomics Facility, NIBB Core Research Facilities, especially Y. Makino, for mass spectrometry analysis, and the Instrument Center, Okazaki, Japan, especially M. Nakano, for HSQC spectra measurements. We also thank M. Yamamoto for experimental assistance, S. Minami for advice on structure similarity comparison and useful discussions, and Y. Ishii and S. Akiyama for valuable comments on the manuscript. The computations were performed using the Research Center for Computational Science (RCCS), Okazaki, Japan.

## Funding

3D-structure determination was supported by Basis for Supporting Innovative Drug Discovery and Life Science Research (BINDS) from AMED under Grant Numbers JP20am0101072 and JP20am0101083. This work was supported by the Japan Society for the Promotion of Science (JSPS) KAKENHI Grants-in-Aid for Scientific Research 15H05592 to N. Koga, 18H05420 to T.K. and N. Koga, 18H05425 to T.M. and 18K06152 to N. Kobayashi, and the Japan Science and Technology Agency (JST) Precursory Research for Embryonic Science and Technology (PRESTO, Grant Number JPMJPR13AD to N. Koga). K. Sakuma was also supported by JSPS KAKENHI Grant-in-Aid for JSPS Research Fellow 15J02427.

## Author contributions

K. Sakuma and N. Koga designed the research. K. Sakuma analyzed natural proteins and performed computational design work. K. Sakuma wrote the program code. K. Sakuma, T.K., and R.K. expressed, purified, characterized the designed proteins by biochemical assay, and prepared protein samples for NMR structure determination: H5_fold-0, H6_fold-C, and H6_fold-Z, by K. Sakuma, and H6_fold-U and H7_fold-K, by T.K. and R.K. For NMR structure determination, Naohiro K., T.S., and T.N. collected data: H5_fold-0_Chantal, H5_fold-C_Rei, and H6_fold-Z_Gogy, by T.S., and H6_fold-U_Nomur and H7_fold-K_Mussoc, by Naohiro K. and T.N. Naohiro K. performed structural analysis. For crystal structure determination, Naoya K. prepared protein samples and K. Suzuki, with advice from T.M., performed crystallization and structural analysis. K. Sakuma, Naohiro K., T.F., K. Suzuki, T.M., T.K., R.K., and N. Koga wrote the manuscript.

## Competing interests

The authors declare no conflicts of interest.

## Data and materials availability

The solution NMR structures of the five designs have been deposited in the Protein Data Bank under the accession numbers 7BQM (H5_fold-0_Chantal), 7BQN (H6_fold-C_Rei), 7BQQ (H6_fold-Z_Gogy), 7BQS (H6_fold-U_Nomur), and 7BQR (H7_fold-K_Mussoc). The NMR data have been deposited in the Biological Magnetic Resonance Data Bank under the accession numbers 36335 (H5_fold-0_Chantal), 36336 (H6_fold-C_Rei), 36337 (H6_fold-Z_Gogy), 36339 (H6_fold-U_Nomur), and 36338 (H7_fold-K_Mussoc). The crystal structure of H5_fold-0_Elsa has been deposited in the Protein Data Bank under the accession number 7DNS.

## Supplementary Materials

Materials and Methods

Supplementary Text

Figures S1-S13

Tables S1-S9

References

## Notes

### Competing Interest Statement

The authors have declared no competing interest.

