## Supplementary Information for "Design of complicated all-α protein structures"

**This PDF file includes:**

Supplementary Text
Figs. S1 to S13
Tables S1 to S11
References

**Supplementary text**

**Detailed description on designed folds**

H5_fold-0

H5_fold-0 is a 5-helix structure (H.O. = 0.22) with 90 residues, of which the backbone structure topology was found as the one with the smallest values for the radius of gyration and H.O. in the combinatorically generated structures. The overall structure shows a snail-like shape with a right-handed screwed α-helices arrangement. This fold comprises four HLH motifs. The first one is the v-shaped type motif (the loop ABEGO pattern is BAABB), similar to the first loop in protein A (PDB code: 1BDD), making the connected helices form a left-handed packing. The second and fourth ones are the corner-type motifs (GBB), identical to one found at the DNA binding site in homeodomains; the loops in the two motifs were found to have different sequences after the Rosetta sequence design, due to different structural environments: GRS and GIT for the second and fourth loops, respectively. The third loop is composed of a residue with the backbone torsion E in the ABEGO representation, making the flanking α-helices v-shaped. The two anti-parallelly aligned α-helices (the first and second helices, and the third and fifth helices) make the space for the protein core, and the shortest fourth helix with eight residues, not making any helix-helix pairings, covers the core space as a lid.

H6_fold-C

H6_fold-C is a 6-helix structure (H.O. = 0.31) with 112 residues. This fold can be divided into two halves, each consisting of anti-parallelly aligned three helices: one is the second, third, and fourth helices, and the other is the first, fifth, and sixth helices. These two halves are arranged crosswise and the core space is formed between the two. This fold comprises five HLH motifs; the first one is the corner-type motif (the loop ABEGO pattern is B); the second one is the hairpin-type motif (GBB) with a left-handed helix-helix packing; the third one is the hairpin-type motif (GB) with a right-handed helix-helix packing; the fourth one is the corner-type motif (B) with a wider bending angle than the first one; the fifth one is the hairpin-type motif (BAAB) with a right-handed helix packing.

H6_fold-Z

H6_fold-Z is a 6-helix structure (H.O. = 0.24) with 121 residues. The five HLH motifs are composed of three corner, one v-shaped, and one hairpin types. This fold shows the Z-like arrangement of the first three helices and the triangle arrangement of three helices from the second to fourth helices. Since the structure contains only one hairpin-type motif, most of the helix-helix packing interactions largely deviate from canonical ones. The fourth HLH motif includes the short GB loop and have a right-handed packing form. The fifth one is the corner type motif (GBB), in which the loop is relatively buried by the third helix, thus one of the hydrogen-bond donors in the loop, the oxygen atom of Gly99, is satisfied through the hydrogen bonding with the Gln49 in the third helix. In addition, this 90° corner bending places the sixth helix in between the first and second helices. The first and sixth helices form a non-local parallel helix-helix packing. The overall topology is complicated and has the arrangement slightly similar to the globin fold.

<u>H6_fold-U</u>

H6_fold-U is a 6-helix structure (H.O. = 0.48) with 113 residues. This fold has an apparent structural similarity to H6_fold-C. This fold also can be divided into two halves, each consisting of anti-parallelly aligned three helices: one is the third, fourth, and fifth helices, and the other is the first, second, and sixth helices. These two halves are arranged almost crosswise and the core space is formed between the two. This fold comprises five HLH motifs; the first one is the hairpin-type motif (the loop ABEGO pattern is BAB); the second one is the v-shaped-type motif (BB); the third one is the hairpin-type motif (AGB); the fourth one is the v-shaped-type motif (AGBB); the fifth one is the corner-type motif (AG). The last HLH motif showed an AGEB loop geometry in the NMR structure, although the overall arrangement of α-helices agreed well. This fold has nonlocal contact patterns similar to Greek-key motifs; non-adjacent secondary structure pairs have many nonlocal contacts.

<u>H7_fold-K</u>

H7_fold-K is a 7 helix structure (H.O. = 0.50) with 128 residues. Among the designed folds, this fold shows the highest H.O. value and bundle-like structure. This fold comprises six HLH motifs. The first HLH motif with a single residue loop (the ABEGO pattern is A) forms a kinked helix structure, which is stabilized by optimized hydrophobic packing around it. The second one is the hairpin-type motif (the loop ABEGO pattern is AGABB); the third one is the hairpin-type motif (AGB); the fourth one is the v-shaped-type motif (AGBB); the fifth one is the v-shaped-type motif (BAB); the sixth one is the corner-type motif (AB). This fold can be divided into two halves: one is the first to the fourth helices and the other is the fifth to the seventh helices. A large hydrophobic core is formed between these two halves, which may explain the highest denaturation temperature among all the five designed folds.

### NMR structure determination

Samples for all designed proteins were stable during NMR structure determination (for more than 2-3 weeks) at the concentration (0.5-1.0 mM), and rarely minor components were found. No noticeable signal changes in 2D $^{1}H$-$^{15}N$ HSQC at different concentrations (2-10 times dilution) were identified.

Structural determination was performed by perfectly blind analysis to avoid any biases being involved in the automated NMR analysis: the analyst never has known the designed structures even their sequences before structure determination. The highly sensitive modern NMR machines (700-800 MHz equipped with the 2nd or 3rd generation of Cryo-probe) and highly concentrated samples yielded a significant number of NOE peaks from NOESY type spectra for all samples. For each sample, around 90% of the NOE peaks were assigned by CYANA and the well-converged structure was obtained, supporting high consensus between the NOE peaks and the calculated NMR structure.

RDC values were exclusively used for assessment of the determined NMR structures. This strategy enhances the reliability of the determined NMR structures after refinement calculations by Amber12, which have the geometrical normality (such as Ramachandran plots and VdW clash) and low violations of restraints. RDC can be affected by local motions of HN-N vectors in a wide range of time scale, but the effect is not large if a sufficient number (the coverage of residues more than 80%) and sufficient amplitude of RDC values ($> \pm 10$ Hz with error $< 0.2$ Hz) are obtained. Moreover, the $\alpha$-helices in the designed structures were designed to have largely different orientations, thus this RDC analysis using $^{1}D_{1H\text{-}15N}$ is suitable for structure validation.

H5_fold-0_Chantal

One of the structural features in the design and determined NMR structures for H5_fold-0_Chantal is the tightly stacked aromatic rings between His63 and Tyr59. The imidazole ring of His63 was assumed to be protonated on Nε2 based on the chemical shift of His63-13Cδ1. Therefore, Nε2 of His63 was protonated in the CYANA and Amber12 calculations. As shown in 2D $^{1}$H-$^{15}$N HSQC in Fig. S5, H5_fold-0_Chantal does not show any minor components at the NMR condition. The signal of Arg22-$^{1}$Hε/$^{15}$Nε is up-field shifted by ring-current effect of Trp33, strongly indicating interactions between side-chains of Arg22 and Trp33. The His63-Hδ2/Cδ2 signal in 2D $^{1}$H-$^{13}$C HSQC supported the tightly packed aromatic rings between His63 and Tyr59. Several methyl and methylene signals are also up-field shifted for Ile74-Hγ2, Val11-Hγ1/2, Leu56-Hγ1/2, Arg22-Hγ2/3 and Arg22-Hβ2/3. These atoms are close to the aromatic rings of Phe14, Tyr18, Trp22, and Tyr59, forming tightly packed hydrophobic core. The structure validation by PALES using almost all of the RDC values (82 out of 88, excluding flexible regions) exhibits the high RMS score, 0.948 (this is higher than designed structure (RMS = 0.894)), indicating that the NMR structure was accurately determined,

<u>H6_fold-C_Rei</u>

The side-chain orientations of the core, Ile, Val, Leu, Met, Phe, and Trp, in the designed structure show remarkable similarity with those of the NMR structure. On the other hand, the side-chain orientation of Trp61 is slightly different between the NMR and designed structures, suggesting that the local packing around the side-chain may not be sufficient enough to fix its conformation. Interestingly, the methyl group of Leu12 located in the position buried by Trp61 does not seem to be shielded. The fluctuation of the Trp61 side-chain likely impairs the packing between Trp61 and Leu12. Only Val76-Hγ1/2, Leu25-Hδ1/2 methyl protons were found to be shielded by Phe82. A lot of methyl groups are considered to contribute the tightly packed hydrophobic core. Interestingly, Gln36 -Hε21 and -Hε22 signals are unusually shielded and down-field shifted. This effect is caused by strong hydrophilic interactions with the side-chains of Gln84 and Glu32, respectively, forming hydrogen bonds to stabilize the helical structure. The side-chain of Asn96 was found to make the N-terminal capping for the following α-helix. The RDC validation score for the determined NMR structure using 102 of 107 RDC values (-24.2~19.83 Hz) except for flexible regions was RMS=0.923 (this is slightly higher than that for the designed protein, RMS=0.916).

H6_fold-Z_Gogy

This protein has a characteristic Met-cluster comprised of the residues 13, 16, 20, 76 and 112. In the NMR structure, the cluster forming a small pocket tightly accommodates the side-chain of Trp73, while in the designed structure, the indole of Trp73 is slightly away from the cluster. Methyl signals of Val72-Hγ1/2, Leu98-Hδ1/2 were up-field shifted by Phe94. Phe94 and Phe108 are packed, supported by the strongly shielded Phe94-Hζ signal by the ring current of the Phe108 side-chain. Although a lot of signals are overlapped in 2D $^{1}H$-$^{15}N$ IPAP-HSQC, $^{1}D_{1H\text{-}15N}$ values (-23.60~30.78 Hz), the signals for 95 of 116 residues, covering all α-helices, were obtained. The RMS value calculated by PALES for the determined structure was 0.913, while that of designed structure was slightly better, 0.921, indicating that both the NMR and designed structures well satisfy experimental data.

H6_fold-U_Nomur

Many amide signal intensities were reduced and some signals were missing because of the buffer condition with higher pH and salt concentration, and some signals were missing. In addition, many amide and methyl signals were crowded in narrow regions because of the high helix content in the structure as well as many Ala residues in the design sequence. The automated assignments for H6_fold-U_Nomur was first performed by FLYA with additional spectra such as 3D HN(CA)CO, HNCA, HN(CO)CA, (H)CC(CO)NH and H(CCCO)NH. The sequence specific assignments were confirmed and corrected by 3D H(CA)NNH, (H)N(CA)NH, HBHA(CO)NH, HNHAHB spectra. The assignments of side-chain signals were also confirmed and corrected by 3D $^{13}C$-HSQC ($^{13}C$-time domain) NOESY $^{13}C$-HSQC, $^{13}C$-HSQC ($^{13}C$-time domain) NOESY $^{15}N$-HSQC and 4D 13C-HSQC NOESY 13C-HSQC spectra. For obtaining RDC values, severely overlapped signals in 2D IPAP $^{1}H$-$^{15}N$ HSQC spectra, 77 of $^{1}D_{1HN\text{-}15N}$ and 64 of $^{1}D_{1Ha\text{-}13Ca}$ were derived from 3D HN(CO)CA-J measured at the same alignment condition. Each value in the RDC table was applied to the PALES calculation, and 141 values were used for the final calculation. The N- and C-terminal helices tend to show larger error in the RDC simulation, indicating that these helices may not be well packed. Despite the low RMS score (0.776) for the designed protein, the overall designed structure should be the same as the NMR structure. On the other hand, the aromatic ring positions of Phe51, Phe54 and Trp43 in the NMR structure were largely different from those in the designed structure. Leu35-Hδ1/2, Leu90-Hδ1/2 are shielded by the ring current, obviously derived from Trp43 and Phe54, respectively. These aromatic rings in the designed structure may not be able to shield any methyl groups of Ile, Val and Leu. These support that the NMR structure is more trustful than the designed structure.

H7_fold-K_Mussoc

Many amide signals were weak or disappeared in $^{1}H$-$^{15}N$ HSQC, because of the higher pH and salt concentration, and many signals were crowded because of the high helix-content in the designed structure. After the automated signal assignments by FLYA, sequence specific assignments were confirmed by 3D (H)N(CA)NH and $^{13}C$-HSQC ($^{13}C$-time domain) NOESY $^{15}N$-HSQC spectra. Side-chain signal assignments for aliphatic residues were corrected by 3D (H)CC(CO)NH, H(CCCO)NH, and (H)CCH-TOCSY. Several proximities of methyl groups were successfully confirmed on 4D NOE spectra to make sure how correctly assigned crowded methyl signals and related NOEs were. The methyl of Leu71-Hδ1/2 are strongly shielded by Phe23 and the methyl group, Ile41-Hγ2, Leu103-Hδ1/2, and Leu113-Hδ1/2, are strongly shielded by Trp109. Trp109-Hζ3 is strongly shielded by the ring current effect of the Phe63 side-chain, indicating the hydrophobic side-chains are correctly packed in the NMR structure.

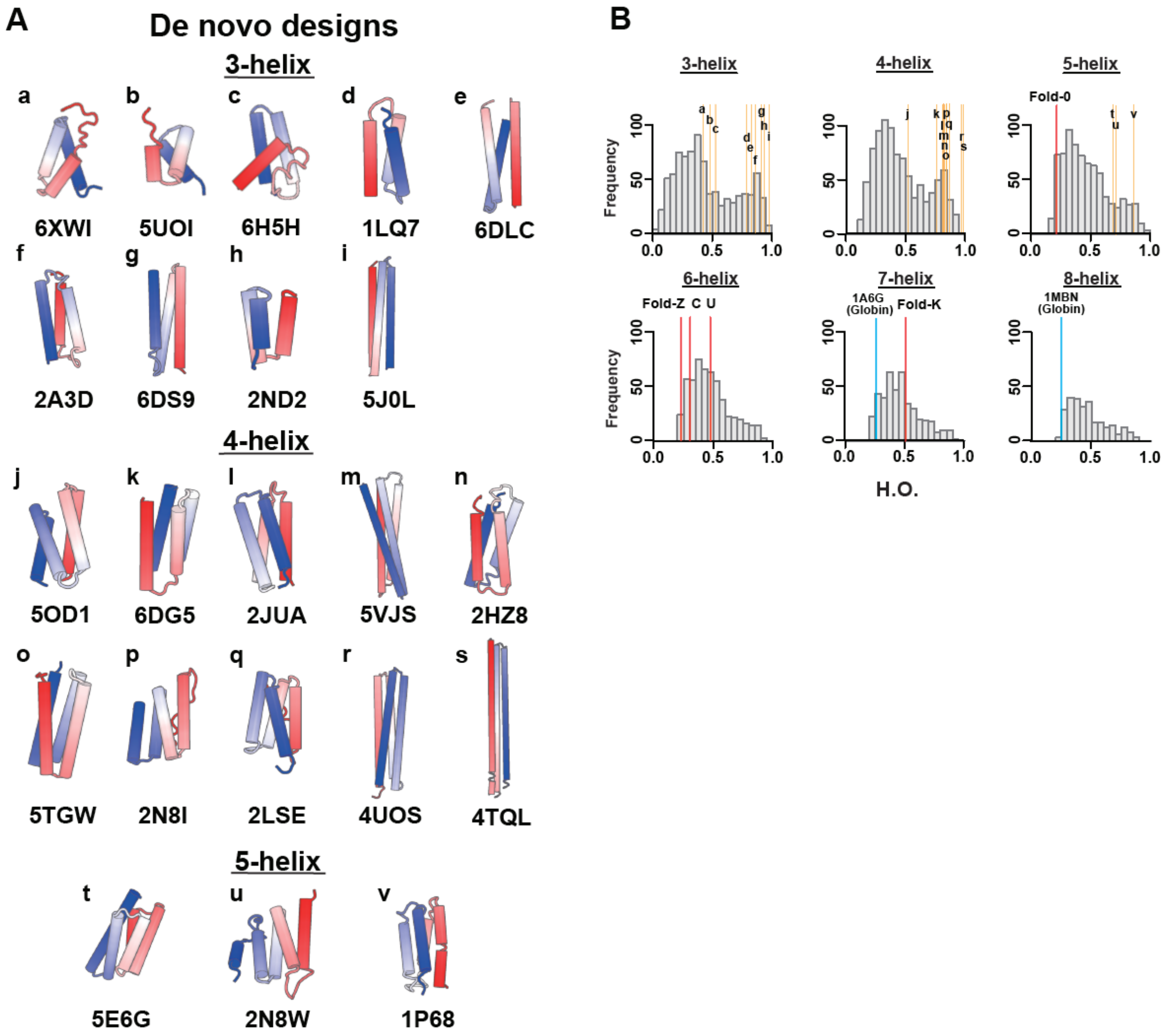

**Fig. S1. 22 de novo designed proteins collected from Protein Data Bank (PDB).**

(A) Structures and their PDB IDs of the 22 de novo designed proteins collected from PDB. (B) The helix order (H.O.) values of the designs were plotted in the H.O. histograms for naturally occurring all-α proteins (these histograms are identical to those in Fig.1).

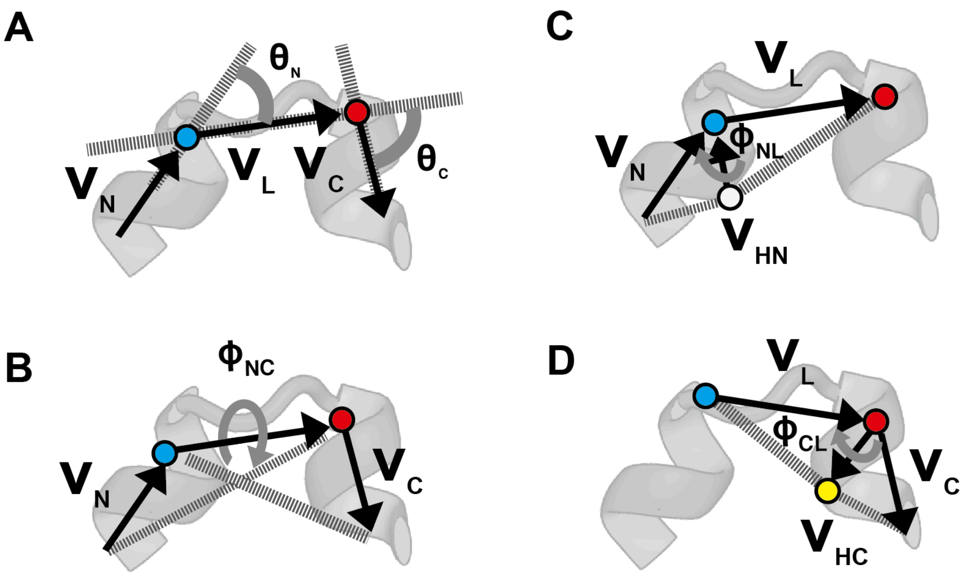

**Fig. S2. The five features representing the HLH tertiary geometry.**

(A-D) For representing tertiary geometries of HLH units, the following angles, $\theta_N$, $\theta_C$, $\phi_{NC}$, $\phi_{NL}$, and $\phi_{CL}$, were identified using the $V_N$, $V_C$, $V_L$, $V_{HN}$, and $V_{HC}$ vectors (these vectors are calculated using Cα atoms). (A) The definitions of $\theta_N$ and $\theta_C$. $V_N$ and $V_C$ respectively represent the helix vectors for the N- and C- terminal helices in a HLH geometry, which are calculated using the equations proposed by Krissinel *et al.* (*18*). $V_L$ is the loop vector from the last Cα atom (blue) in the N-terminal helix to the first Cα atom (red) in the C-terminal helix. $\theta_N$ was identified as the angle between the $V_N$ and $V_L$ vectors; $\theta_C$, was identified as the angle between the $V_C$ and $V_L$ vectors. (B) The definitions of $\phi_{NC}$. $\phi_{NC}$ was identified as the dihedral angle between the plane defined with the $V_N$ and $V_L$ vectors and that with the $V_C$ and $V_L$ vectors. (C) The definition of $\phi_{NL}$. $V_{HN}$ is the helix spiral vector at the end of the N-terminal helix, which was identified as the vector pointed to the last Cα atom (blue) in the N-terminal helix from the Cα atom immediately before the last Cα atom (white). $\phi_{NL}$ was identified as the dihedral angle between the plane defined with the $V_{HN}$ and $V_L$ vectors and that with the $V_{HN}$ and $V_N$ vectors. (D) The definition of $\phi_{CL}$. $V_{HC}$ is the helix spiral vector at the beginning of the C-terminal helix, which was identified as the vector from the first Cα atom (red) in the C-terminal helix to the Cα atom immediately after the first Cα atom (yellow). $\phi_{CL}$ was identified as the dihedral angle between the plane defined with the $V_C$ and $V_{HC}$ vectors and that with the $V_L$ and $V_{HC}$ vectors.

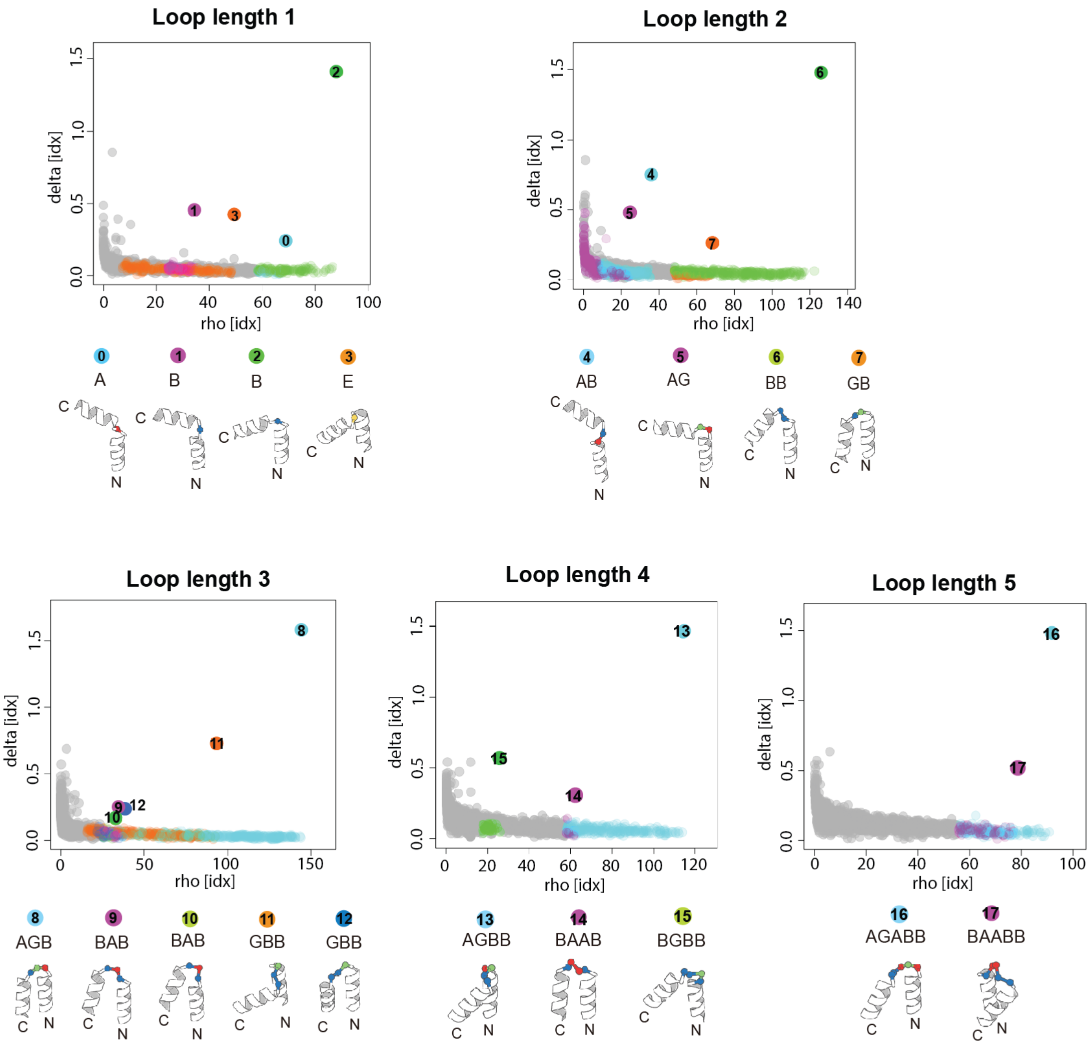

**Fig. S3. Statistical analysis of HLH motifs of naturally occurring proteins.**

HLH structures collected from naturally occurring protein structures were clustered for each loop length from one to five based on the pairwise Euclidean distance between the five-dimensional vectors of the features shown in fig. S2, using the density clustering algorithm (*4*). For each loop length, decision graphs to determine density peaks of clusters are shown, in which rho represents the local density of a point in the five-dimensional feature vector space and delta represents the minimum distance between a point to any other point with higher density; for the point with highest density, delta is calculated as the maximum distance to any other points.

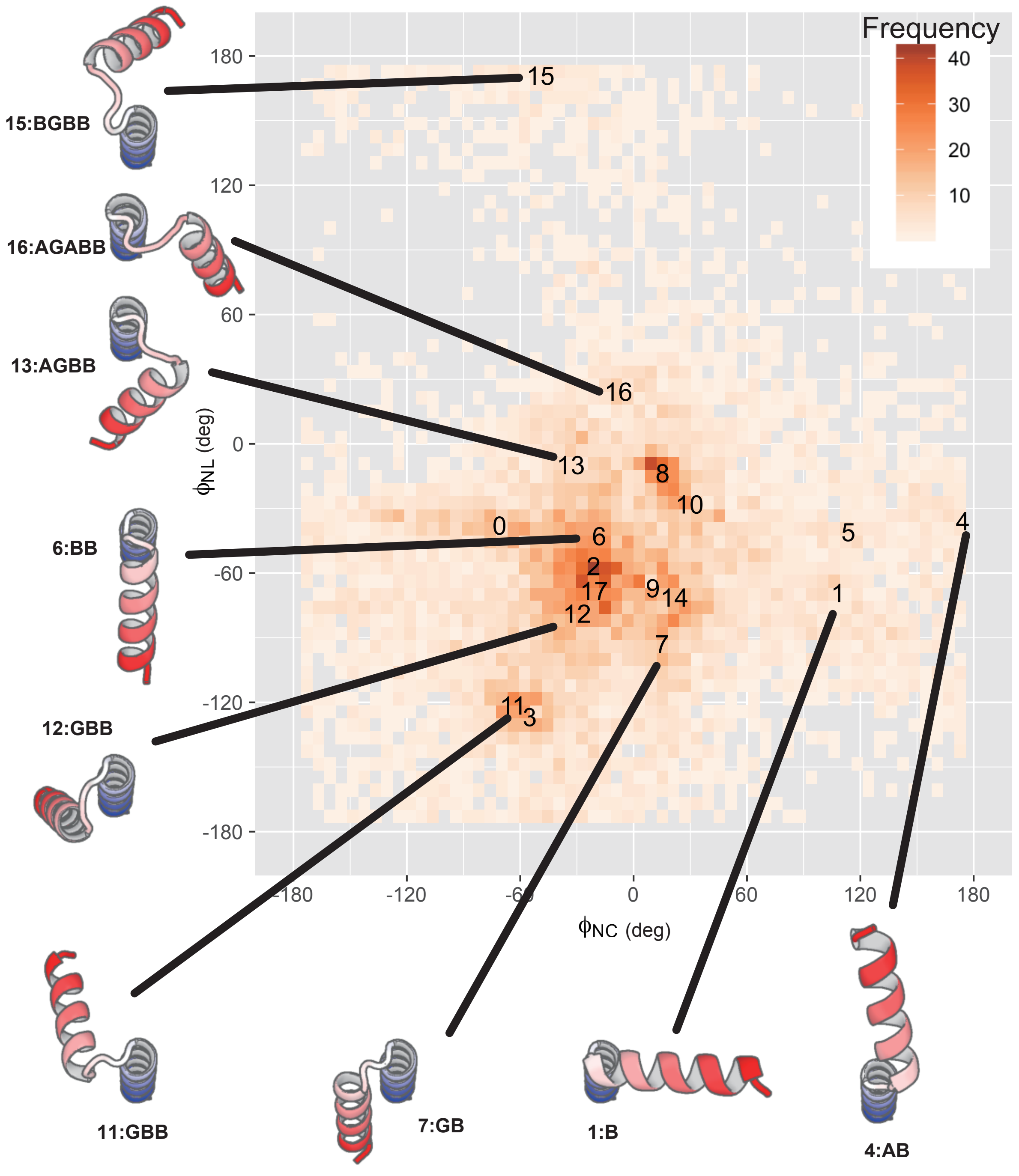

**Fig. S4. Mapping of 18 representative HLH motifs on the $\phi_{NC}$-$\phi_{NL}$ plane.**

The loop numbers with their ABEGO torsion patterns correspond to the 18 representative HLH motifs shown in fig. S3.

**Fig. S5. ABEGO-based loop geometries and amino acid sequence preferences of the cluster that each HLH motif belongs to.**

(Left) The 18 representative HLH motifs are shown as in fig. S3. (Middle) ABEGO torsion patterns of the loop. This result suggests that the relative arrangements of adjacent helices strongly limit the torsion patterns of the connecting loop. (Right) Amino acid sequence preferences of each HLH motif. The first residue of the loop is indicated by an arrow.

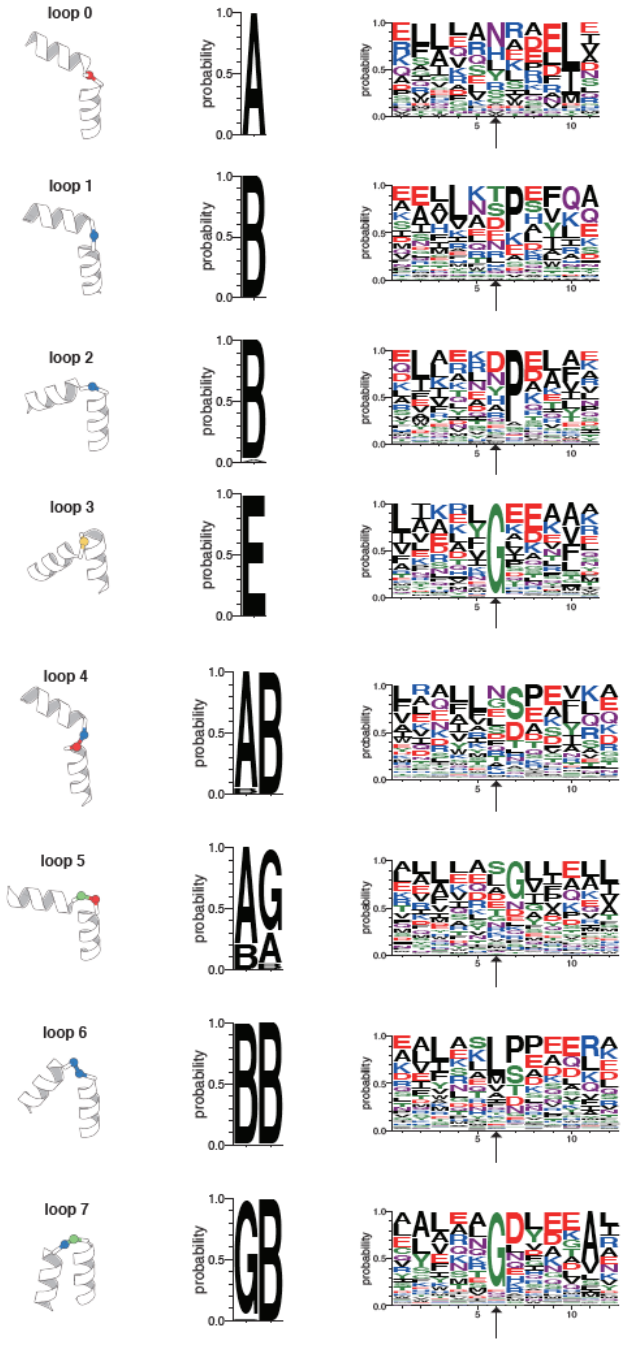

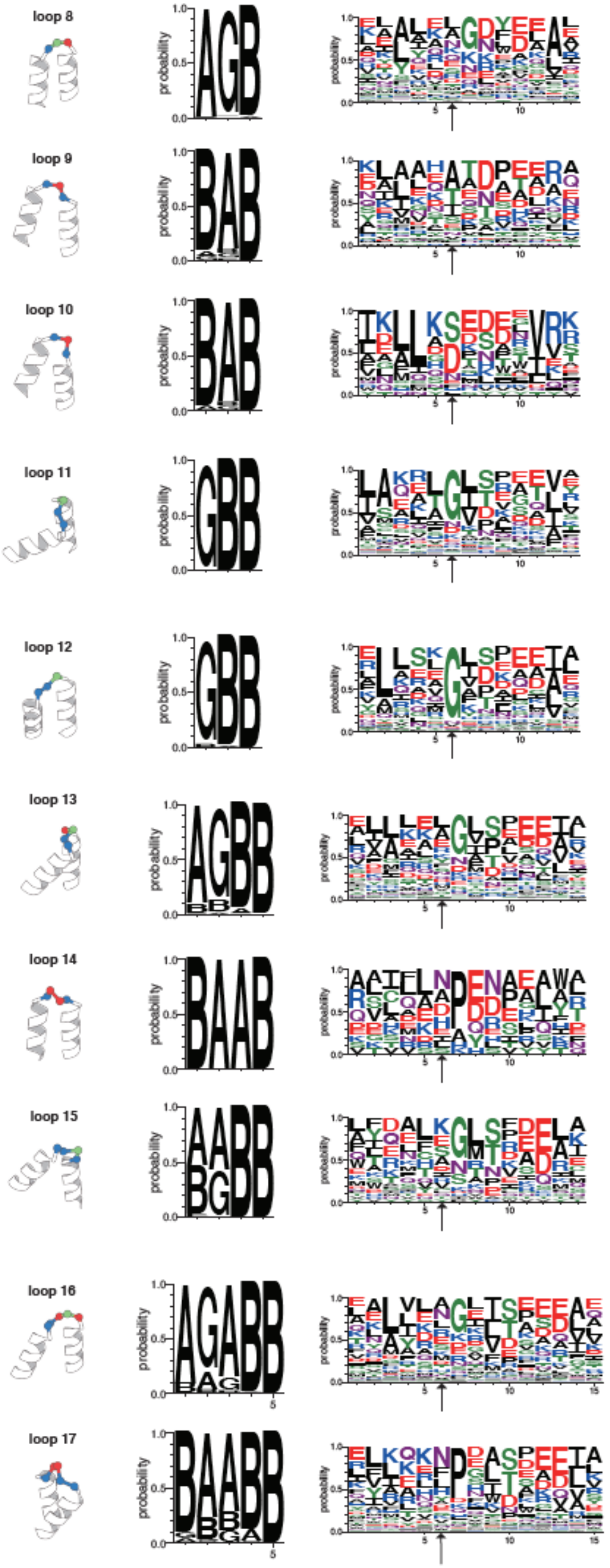
loop 8
loop 9
loop 10
loop 11
loop 12
loop 13
loop 14
loop 15
loop 16
loop 17

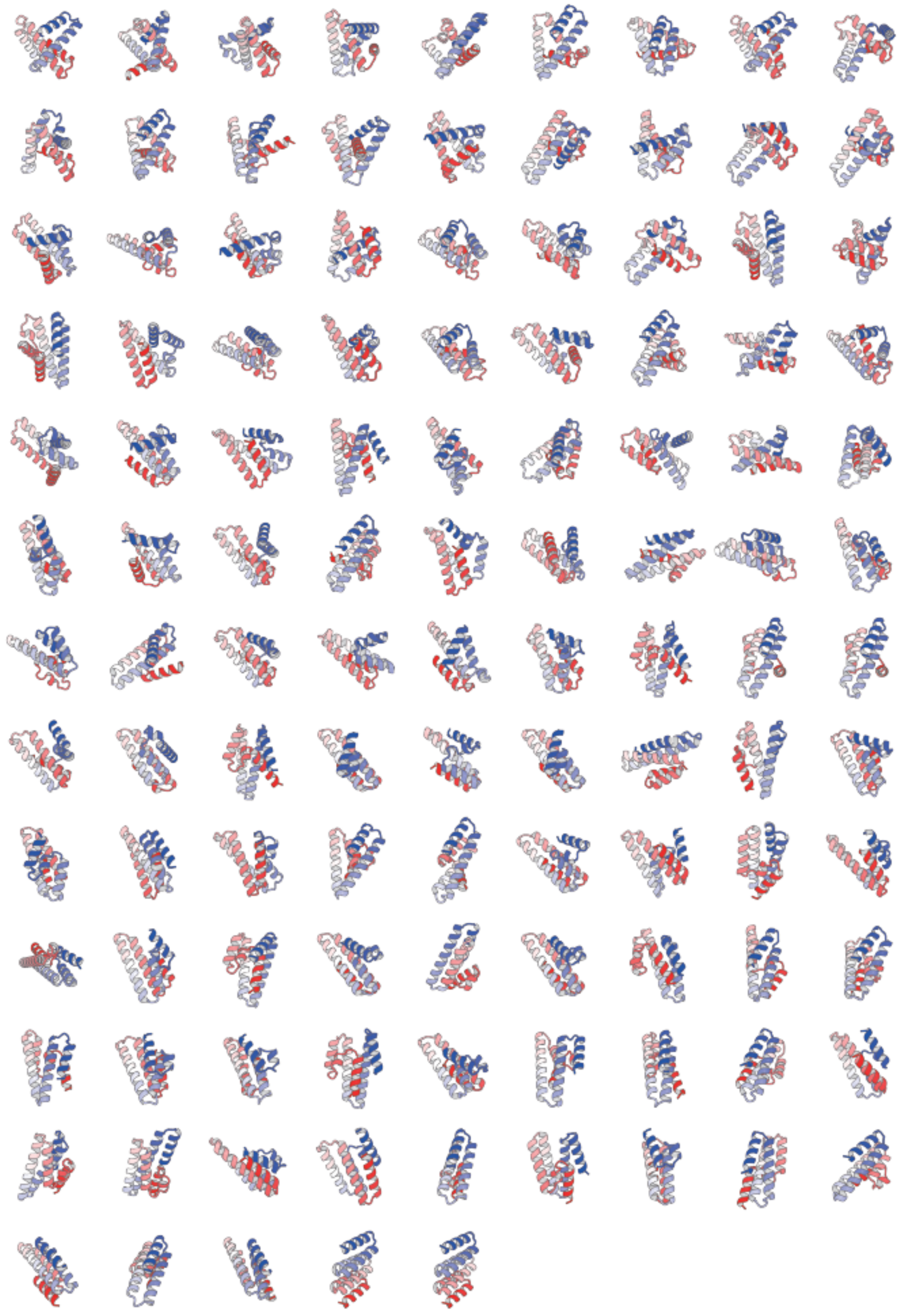

**Fig. S6. Examples of compact structures obtained from the enumeration of 6-helix structures.**

The structures are sorted by their H.O. values from top left to bottom right. The top left structure shows the smallest H.O. value and has irregularly packed α-helices, whereas the bottom right one shows the highest H.O. value and has parallelly aligned α-helices.

**Fig. S7. Design backbone structures with ABEGO loop types.**

Designed backbone structures with different views to clearly show the details around each loop. The residues with the backbone torsion angle, A, B, E, and G, in the ABEGO representation, are shown in red, blue, yellow, and green, respectively.

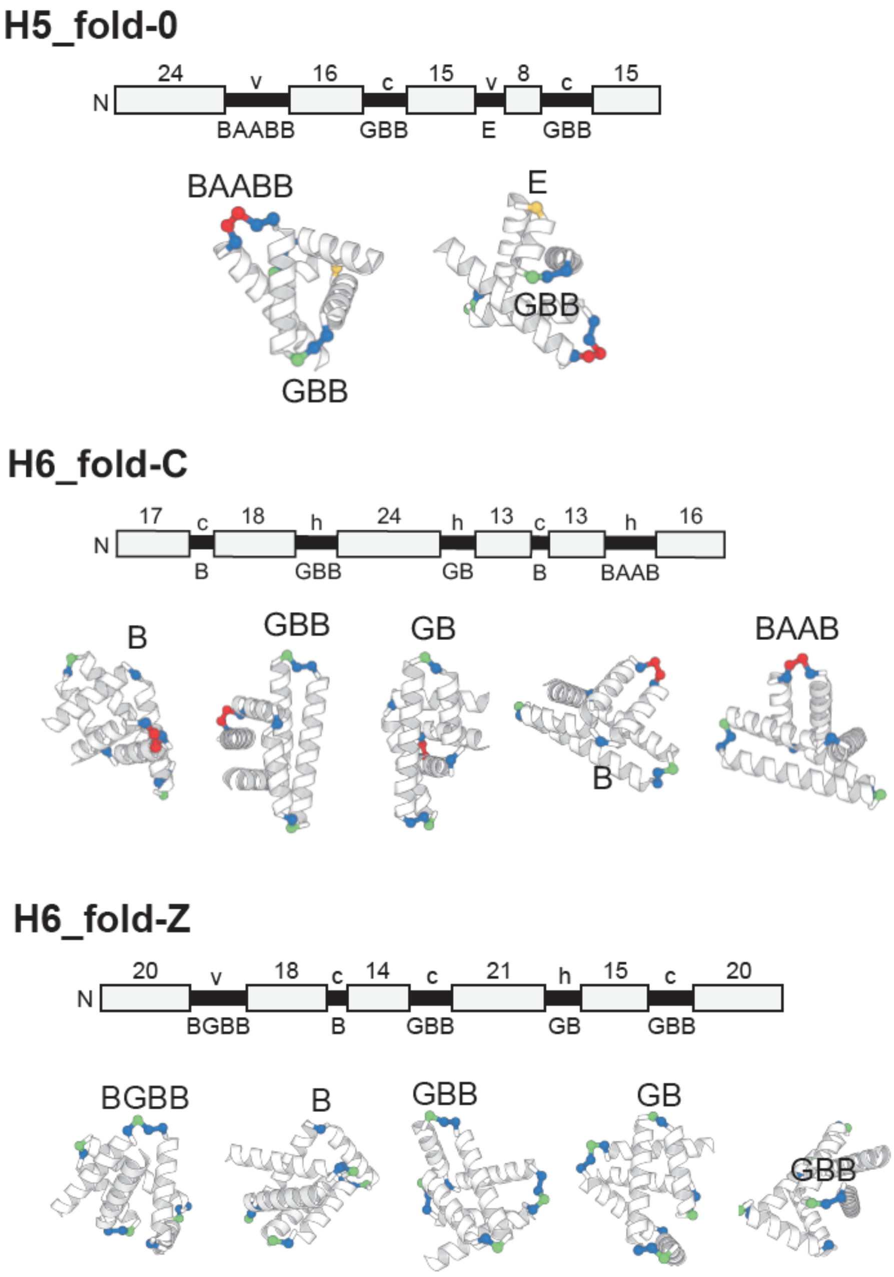

### H6_fold-U

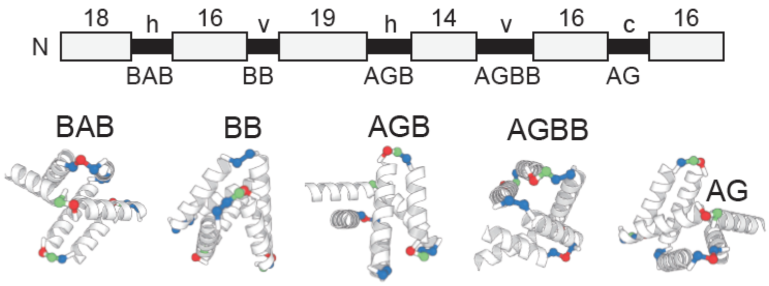

### H7_fold-K

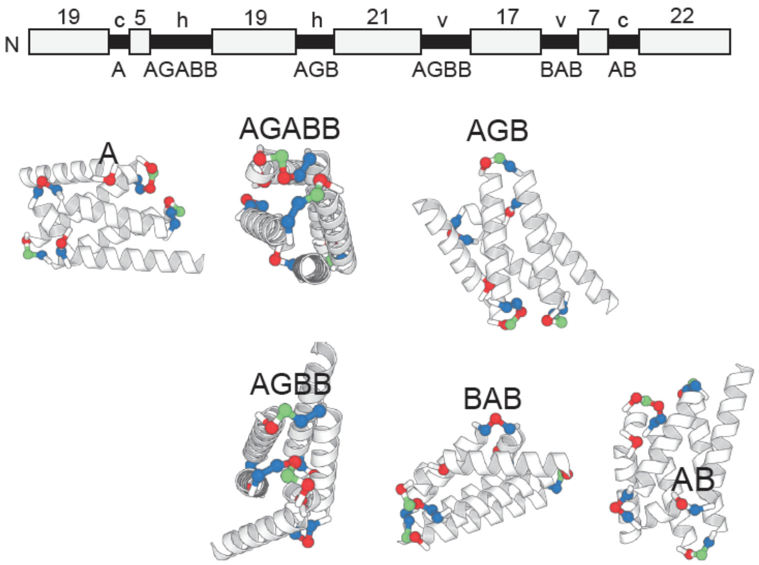

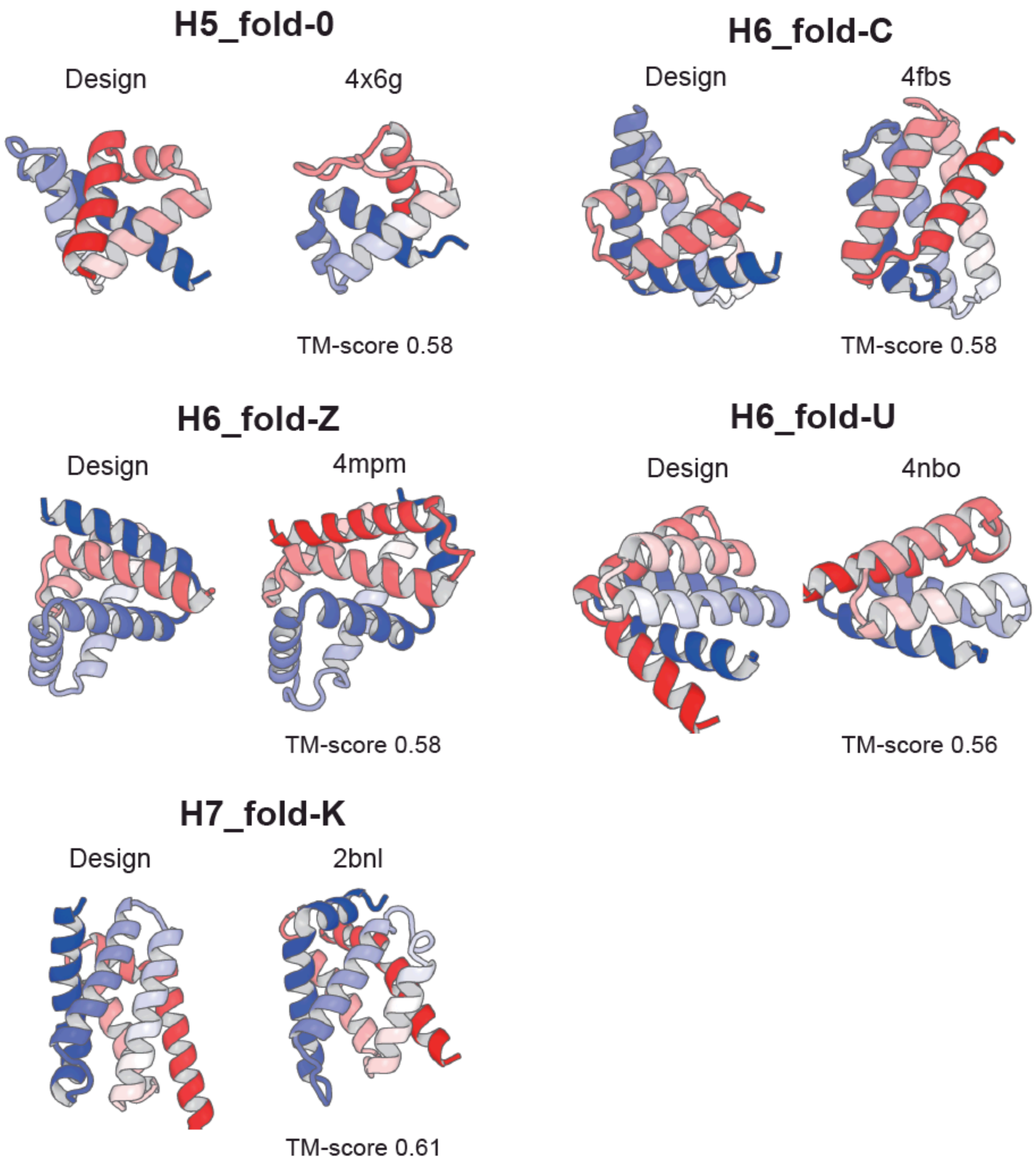

**Fig. S8. Comparison of designed structures and the most similar naturally occurring proteins.**

The designed structures (left) and the most similar ones (right) with pdb ids and TM-score values.

A

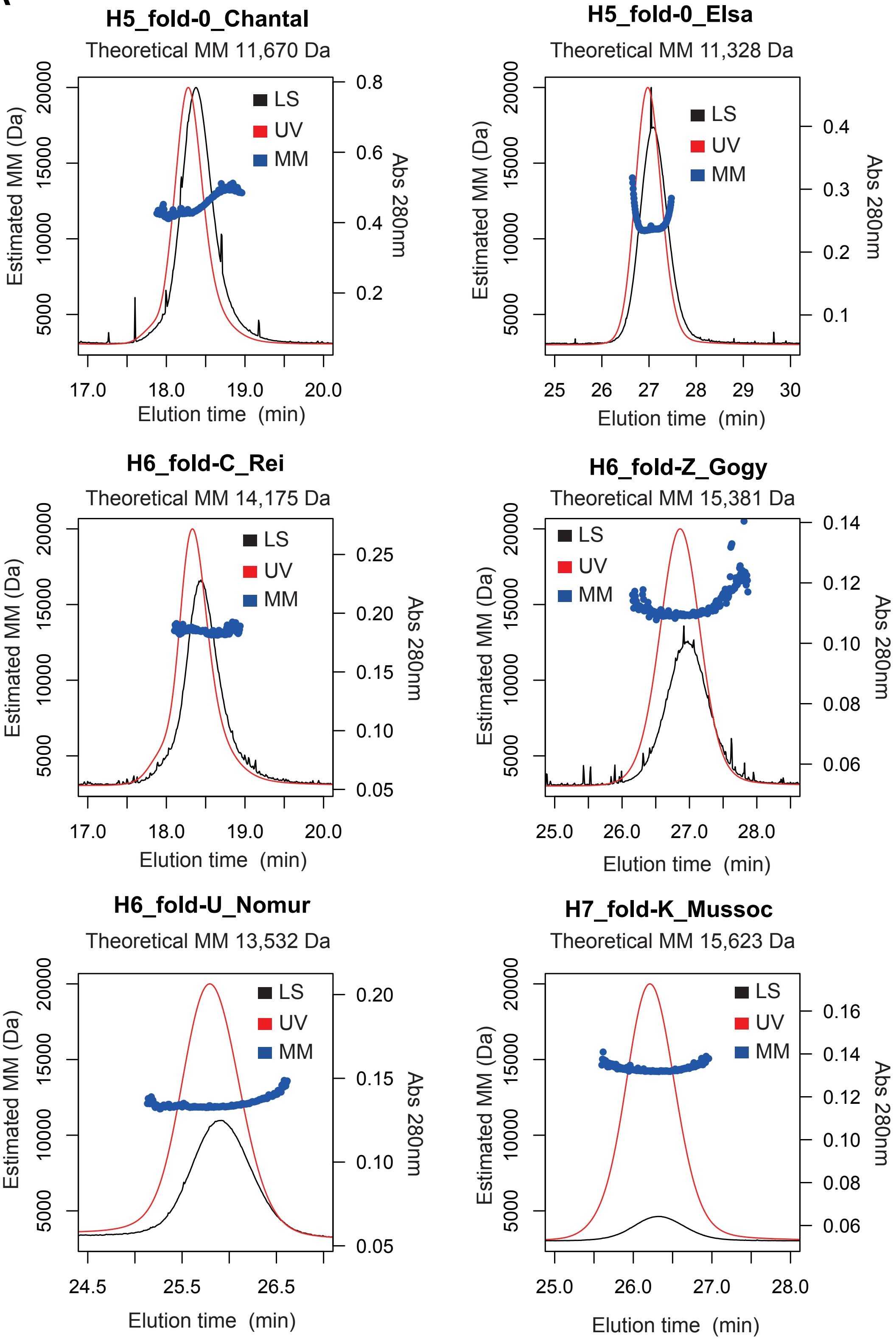
H5_fold-0_Chantal
Theoretical MM 11,670 Da
LS
UV
MM
Estimated MM (Da)
Abs 280nm
Elution time (min)
17.0
18.0
19.0
20.0
5000
10000
15000
20000
0.2
0.4
0.6
0.8
H5_fold-0_Elsa
Theoretical MM 11,328 Da
25
26
27
28
29
30
0.1
0.2
0.3
0.4
H6_fold-C_Rei
Theoretical MM 14,175 Da
0.05
0.10
0.15
0.20
0.25
H6_fold-Z_Gogy
Theoretical MM 15,381 Da
25.0
26.0
27.0
28.0
0.06
0.08
0.10
0.12
0.14
H6_fold-U_Nomur
Theoretical MM 13,532 Da
24.5
25.5
26.5
0.05
0.10
0.15
0.20
H7_fold-K_Mussoc
Theoretical MM 15,623 Da
0.06
0.08
0.10
0.12
0.14
0.16

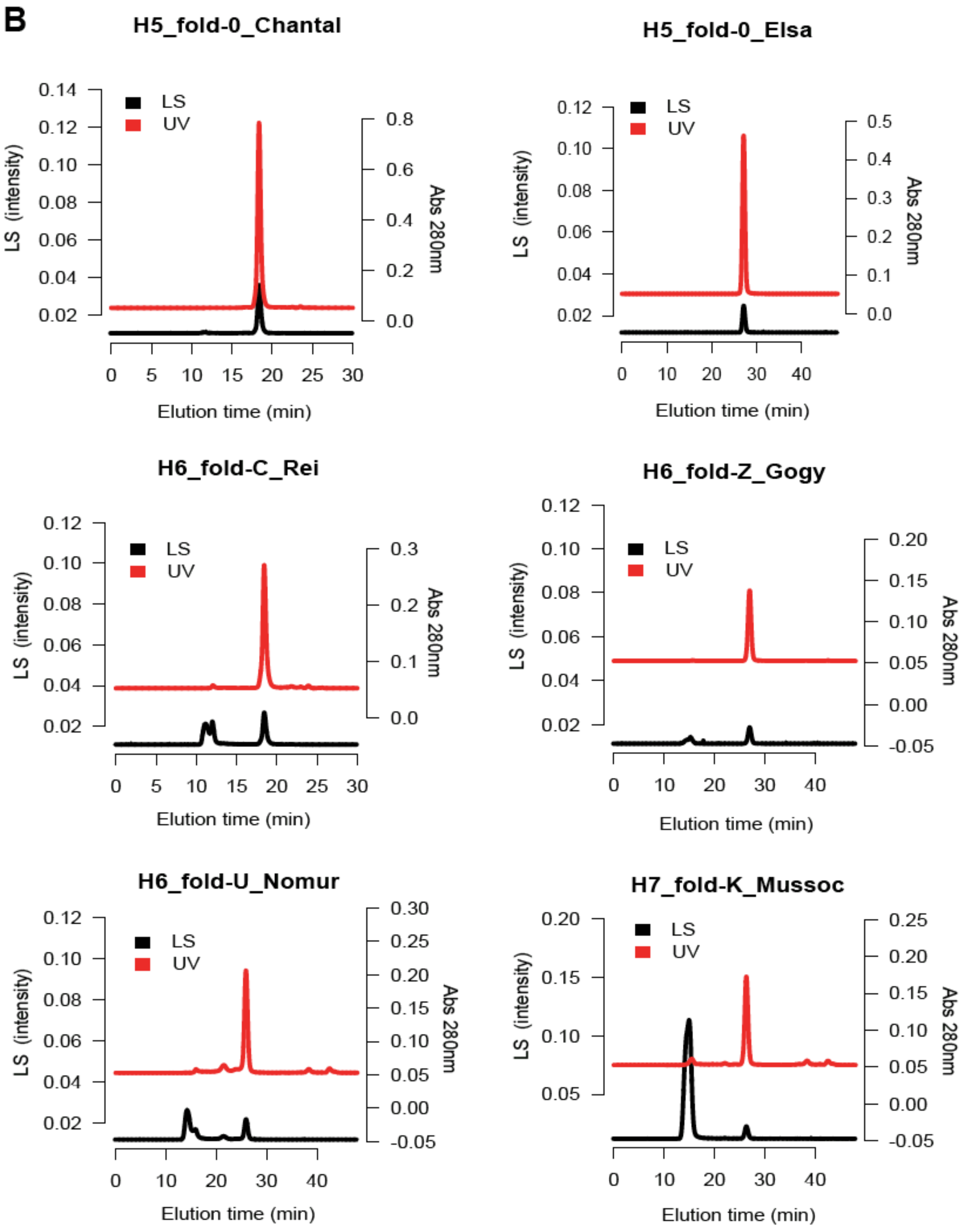

**Fig. S9. Results of SEC-MALS.**

**(A)** Close-up views around the dominant peaks. **(B)** SEC chromatograms of nickel-purified designed proteins. LS: light scattering, UV: ultraviolet, MM: molecular mass.

**Fig. S10. 2D 1H-15N HSQC spectra of experimentally determined structures.**

Assigned peaks are labeled with residue numbers. For side-chain amide atoms, the labels are accompanied with d or e, representing Hδ1/Nδ and Hδ2/Nδ for Asn, and Hε1/Nε and Hε2/Nε for Gln, respectively.

#### H5_fold-0-Chantal

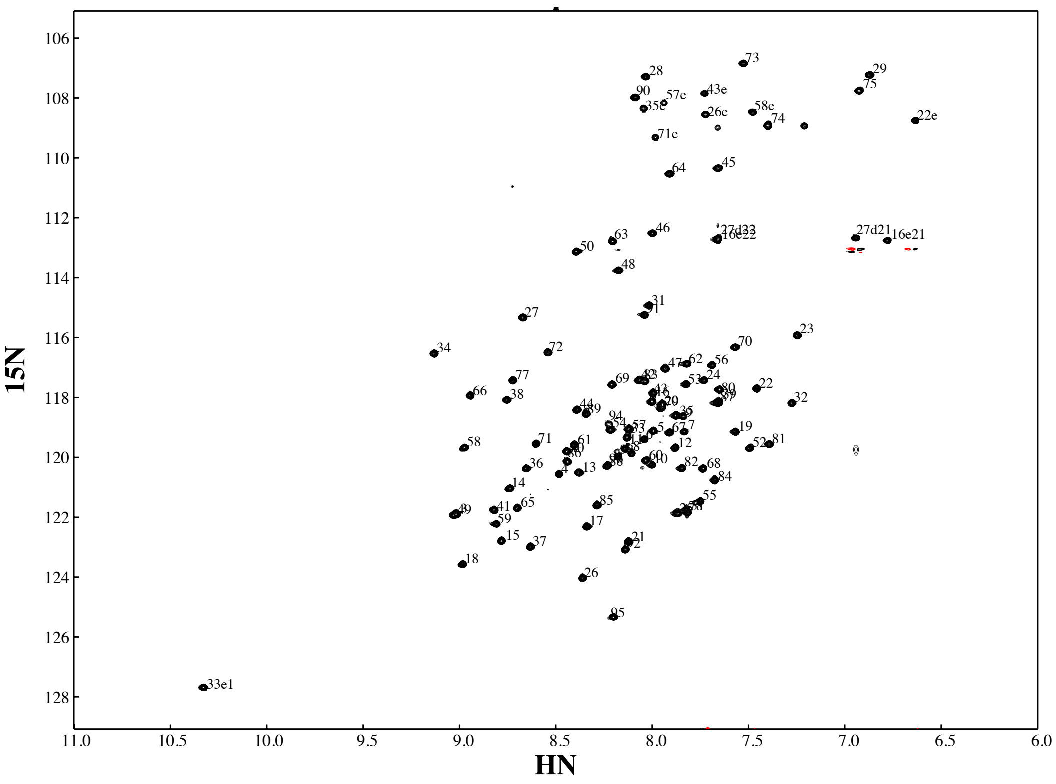

### H6_fold-C_Rei

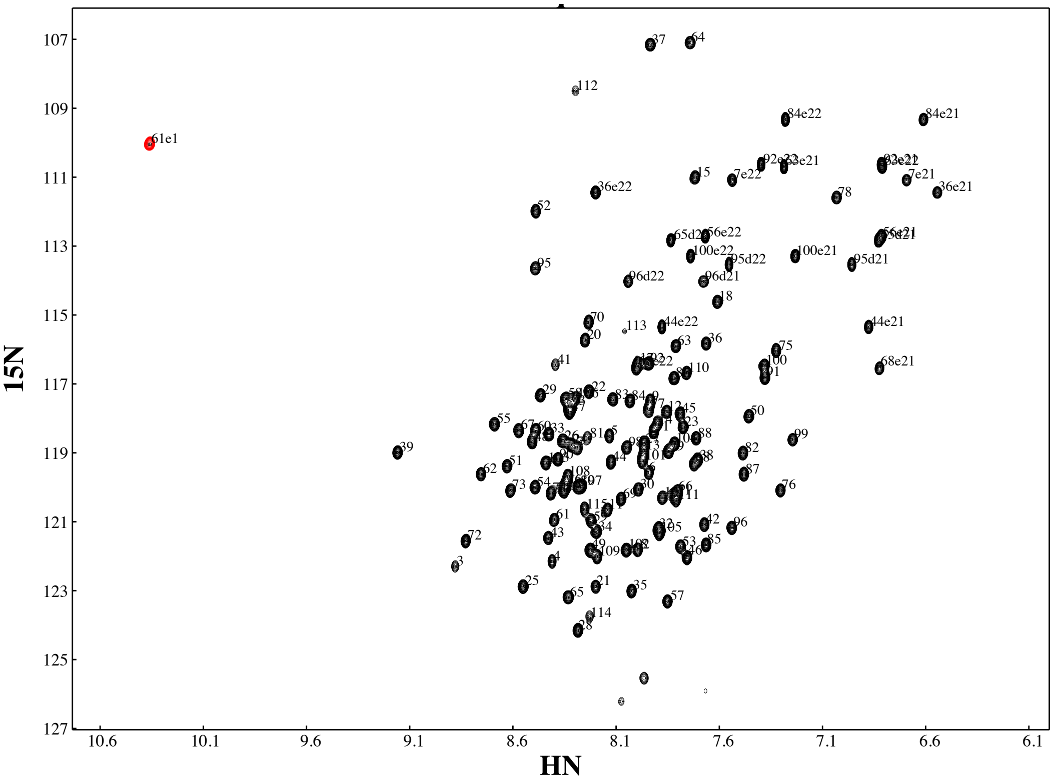

### H6_fold-Z_Gogy

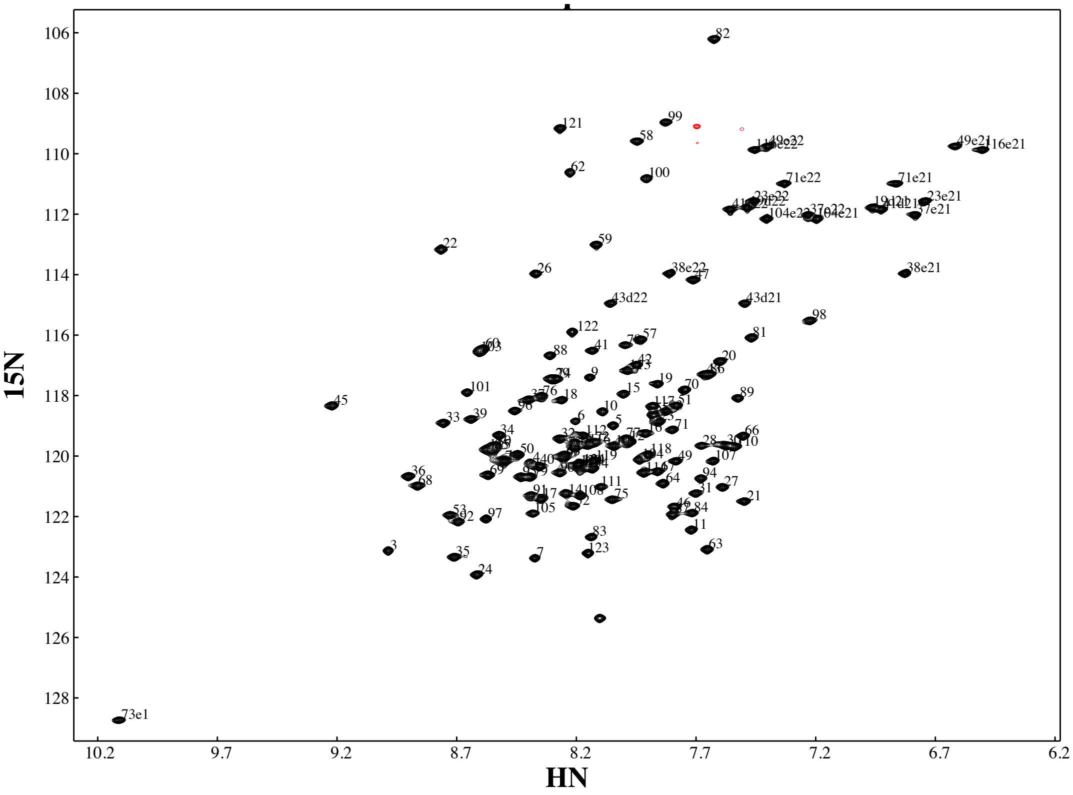

### H6_fold-U_Nomur

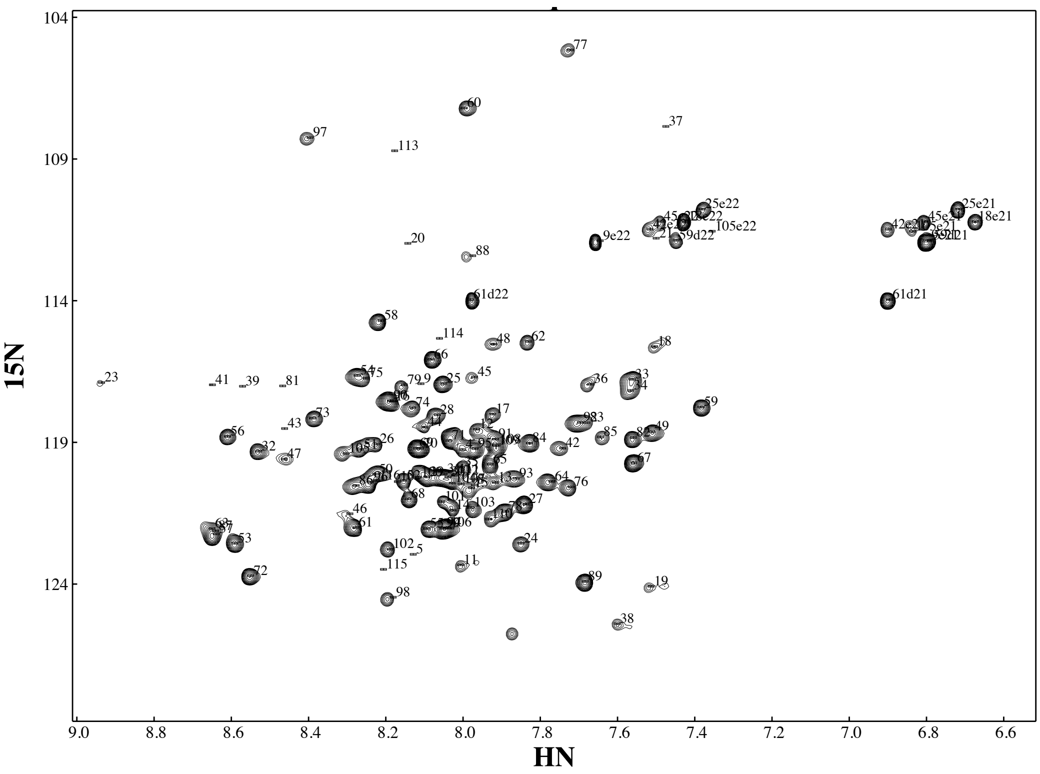

### H7_fold-K_Mussoc

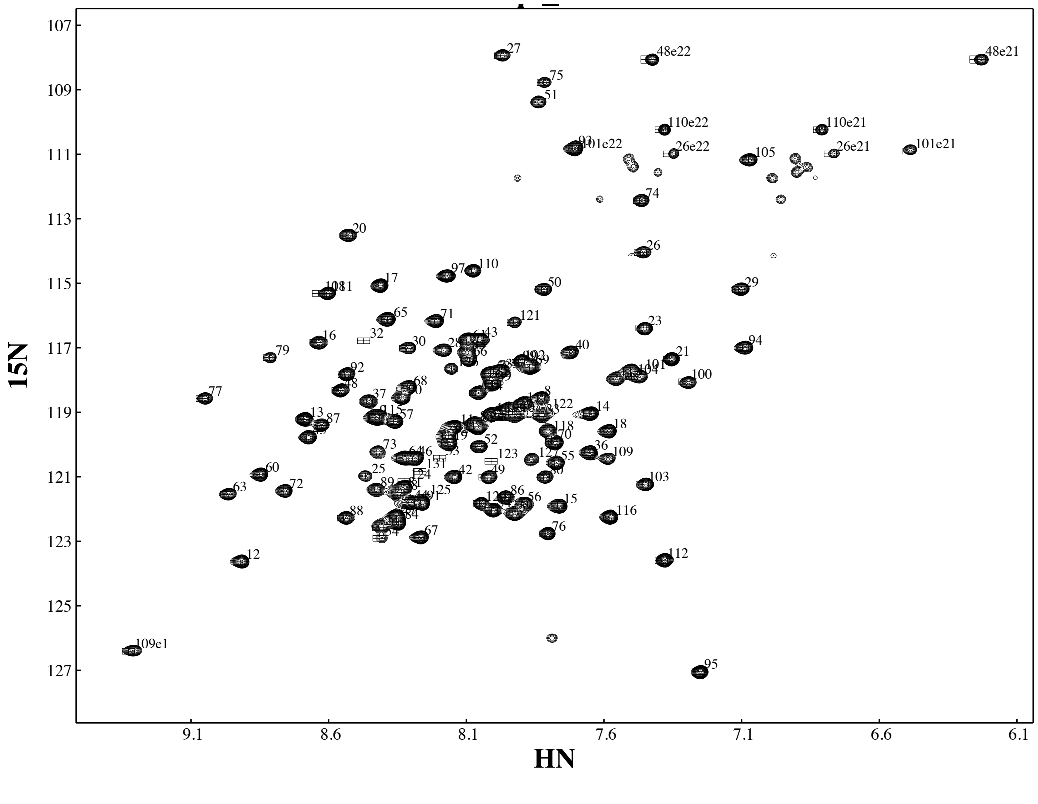

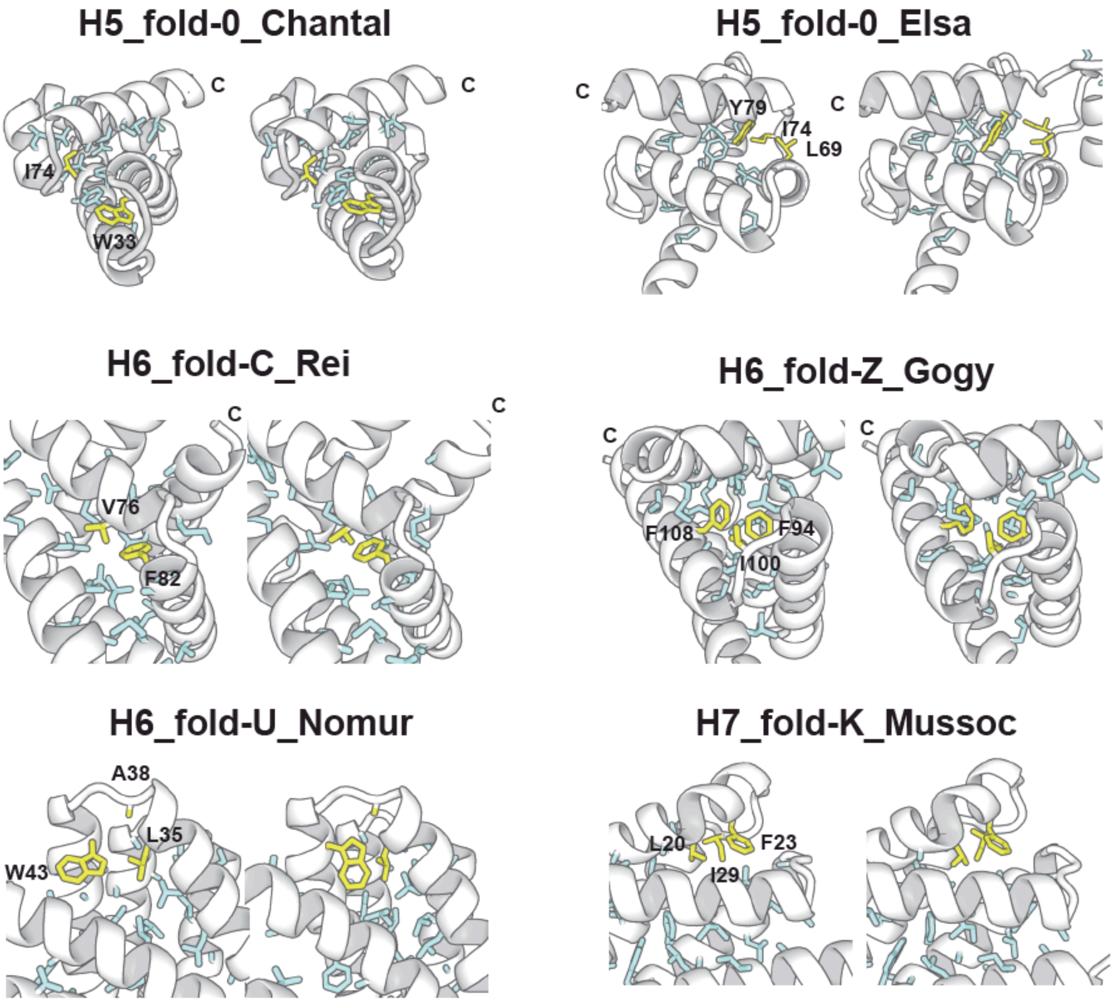

**Fig. S11. Comparison of computational models with experimentally determined structures.** Hydrophobic core residues are shown in stick. Balky hydrophobic side-chains from loops and the neighboring α-helices, which spiked the core and pinned the loops to the target conformations, are shown in yellow.

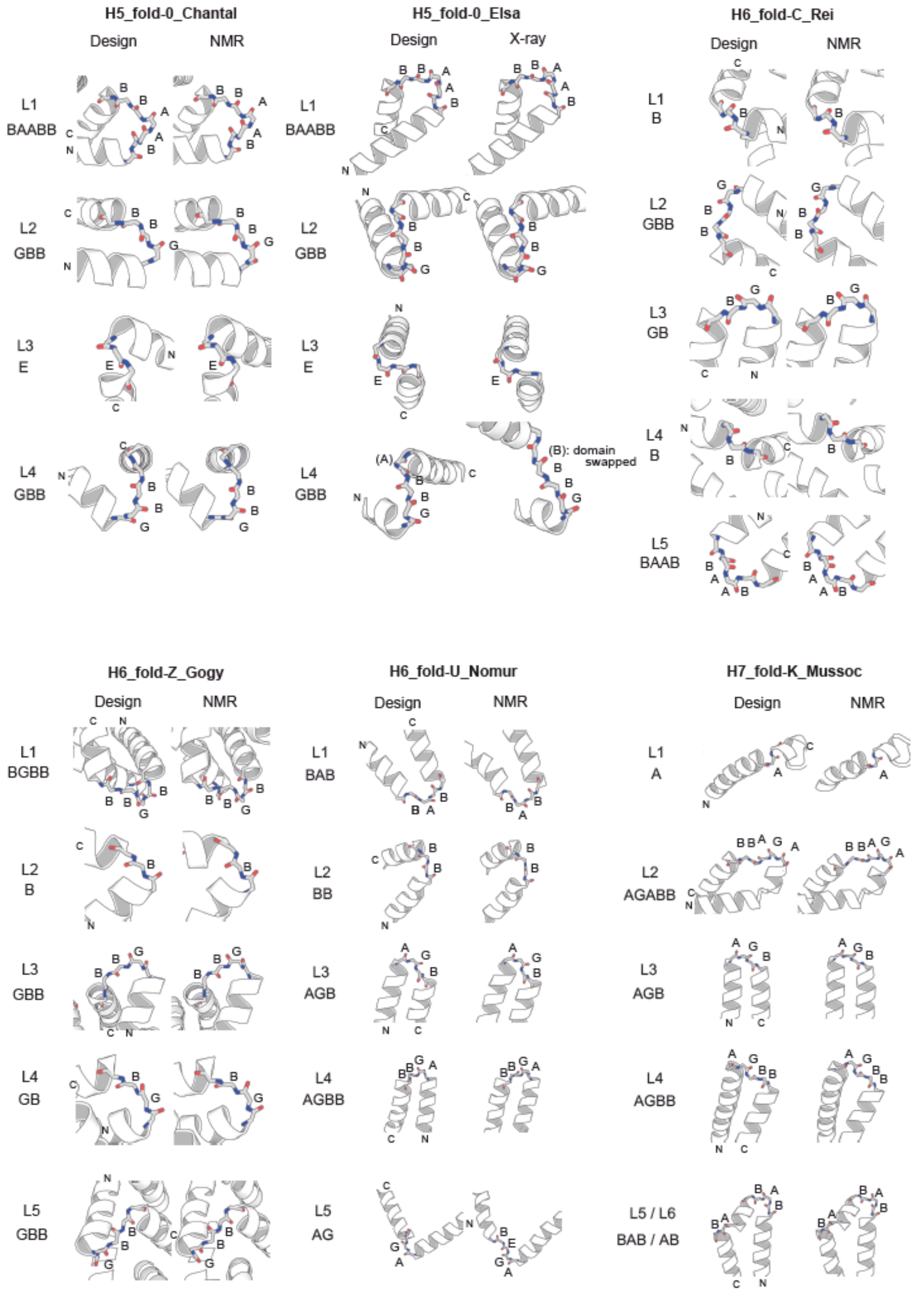

**Fig. S12. Comparison of loop geometries of HLH motifs between design models and experimental structures.**

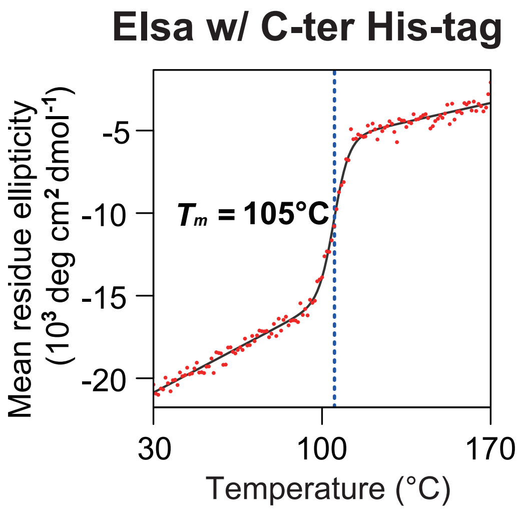

**Fig. S13. Thermal denaturation of Elsa with the C-terminal His-tag.**

Thermal denaturation was measured at 222 nm by CD. The data were fit to a two-state model (solid line) to obtain the $T_m$.

| | #designs tested | Expressed[1] | Soluble[1] | α-protein CD spectrum (20 °C) | Monomeric[2] | Well resolved NMR[3] | Success (rate %) |
|---|---|---|---|---|---|---|---|
| **H5_fold-0** | 10 | 10 | 9 | 9 | 7 | 5 | 5 (50) |
| **H6_fold-C** | 7 | 7 | 6 | 6 | 5 | 3 | 3 (43) |
| **H6_fold-Z** | 7 | 7 | 7 | 7 | 4 | 4 | 4 (57) |
| **H6_fold-U** | 8 | 8 | 5 | 5 | 5 | 4 | 4 (50) |
| **H7_fold-K** | 8 | 8 | 7 | 7 | 6 | 6 | 6 (75) |

**Table S1. Summary of experimental results for designed proteins.**

The second column shows the number of designs experimentally tested for the fold in the leftmost column. The subsequent columns give the number of designs that satisfy each experimental characterization, which was performed sequentially from the left to the right. The successful designs are defined as those that satisfy all criteria and are expected to fold into correct fold. The details of the results are shown in tables S2-6.

[1] Expression and solubility were assessed by SDS-PAGE and mass spectrometry.

[2] SEC-MALS was used to determine oligomerization state. The number of designs in which the main peak of the absorbance at 280 nm corresponds to the monomeric state was counted.

[3] $^{1}H$-$^{15}N$ HSQC spectra were collected.

| | Expressed | Soluble | α-protein CD spectrum (20 °C) | Monomeric | Well-resolved HSQC |
|---|---|---|---|---|---|
| **Ana** | Y | N | | | |
| **Bill** | Y | Y | Y | Y | Y |
| **Chantal** | Y | Y | Y | Y | Y |
| **Chris** | Y | Y | Y | N | |
| **Danny** | Y | Y | Y | N | |
| **Elsa** | Y | Y | Y | Y | Y |
| **Fay** | Y | Y | Y | Y | † |
| **Franklin** | Y | Y | Y | Y | Y |
| **Hermine** | Y | Y | Y | Y | N |
| **Lary** | Y | Y | Y | Y | Y |

**Table S2. Summary of experimental results of 10 designs for H5_fold-0.**

Each row corresponds to the results for each design. The columns give the results for each experimental test, of which the details are described in table S1. Each test was performed sequentially from the left to the right; well-behaved designs at one test (Y) go to the next one and not well-behaved designs (N) end the tests.

† The experiment was not conducted due to low concentration.

| | Expressed | Soluble | α-protein CD spectrum (20 °C) | Monomeric | Well-resolved HSQC |
|---|---|---|---|---|---|
| **Cyrus** | Y | Y | Y | Y | Y |
| **Dan** | Y | Y | Y | Y | Y |
| **Max** | Y | Y | Y | Y | ‡ |
| **Melvin** | Y | N | | | |
| **Moca** | Y | Y | Y | Y | † |
| **Paul** | Y | Y | Y | N | |
| **Rei** | Y | Y | Y | Y | Y |

**Table S3. Summary of experimental results of 7 designs for H6_fold-C.**

The summary was given in the same way as table S2.

‡ The experiment was not conducted due to high nucleotide concentration.

† The experiment was not conducted due to low concentration.

| | Expressed | Soluble | α-protein CD spectrum (20 °C) | Monomeric | Well-resolved HSQC |
|---|---|---|---|---|---|
| **Gogy** | Y | Y | Y | Y | Y |
| **Hoto** | Y | Y | Y | N | |
| **Kome** | Y | Y | Y | N | |
| **Nazu** | Y | Y | Y | Y | Y |
| **Seri** | Y | Y | Y | Y | Y |
| **Siro** | Y | Y | Y | Y | Y |
| **Suzu** | Y | Y | Y | N | |

**Table S4. Summary of experimental results of 7 designs for H6_fold-Z.**

The summary was given in the same way as table S2.

| | Expressed | Soluble | α-protein CD spectrum (20 °C) | Monomeric | Well-resolved HSQC |
|---|---|---|---|---|---|
| **Cacyd** | Y | Y | Y | Y | ‡ |
| **Hatake** | Y | Y | Y | Y | Y |
| **Kazi** | Y | N | | | |
| **Morit** | Y | N | | | |
| **Nomur** | Y | Y | Y | Y | Y |
| **Sait** | Y | N | | | |
| **Sho** | Y | Y | Y | Y | Y |
| **Takin** | Y | Y | Y | Y | Y |

**Table S5. Summary of experimental results of 8 designs for H6_fold-U.**

The summary was given in the same way as table S2.

‡ The experiment was not conducted due to high nucleotide concentration.

| | Expressed | Soluble | α-protein CD spectrum (20 °C) | Monomeric | Well-resolved HSQC |
|---|---|---|---|---|---|
| **Chario** | Y | N | | | |
| **Dark** | Y | Y | Y | N | |
| **Jazzbird** | Y | Y | Y | Y | Y |
| **Mussoc** | Y | Y | Y | Y | Y |
| **NewWave** | Y | Y | Y | Y | Y |
| **R3** | Y | Y | Y | Y | Y |
| **Rush** | Y | Y | Y | Y | Y |
| **Third** | Y | Y | Y | Y | Y |

**Table S6. Summary of experimental results of 8 designs for H7_fold-K.**

The summary was given in the same way as table S2.

| Design ID | H5_fold-0 _Chantal | H6_fold-C _Rei | H6_fold-Z _Gogy |
|---|---|---|---|
| PDB ID | 7BQM | 7BQN | 7BQQ |
| BMRB Entry | 36335 | 36336 | 36337 |
| **NMR distance and dihedral constraints** | | | |
| **Detected NOE peaks** | | | |
| $^{15}$N-edited NOESY | 1,308 | 2,141 | 2,074 |
| $^{13}$C-edited NOESY for aliphatic | 3,847 | 3,986 | 3,398 |
| $^{13}$C-edited NOESY for aromatic | 217 | 103 | 239 |
| **Distance constraints** | | | |
| Total | 2,098 (100.0%) | 3,010 (100.0%) | 2,771 (100.0%) |
| Intra-residue | 425 (20.3%) | 596 (19.7%) | 551 (19.9%) |
| Inter-residue | | | |
| Sequential (\|i-j\| = 1) | 555 (26.5%) | 727 (24.1%) | 620 (22.4%) |
| Medium-range (1 < \|i-j\| < 5) | 617 (29.4%) | 983 (32.78%) | 867 (31.3%) |
| Long-range (\|i-j\| ≥ 5) | 501 (23.9%) | 712 (23.6%) | 733 (26.5%) |
| **Dihedral constraints** | | | |
| Total | 132 | 206 | 232 |
| **NMR structure statistics** | | | |
| **Target function (Å$^2$)** | | | |
| Total | 1.10±0.09 | 1.12±0.007 | 2.62±0.22 |
| Upl¶ | 0.0055±0.0009 | 0.0028±0.0062 | 0.0065±0.0010 |
| VDW | 5.3±0.03 | 5.8±0.30 | 9.6±0.7 |
| Dihedral | 0.68±0.04 | 0.52±0.03 | 0.76±0.07 |
| **Amber structure statistics (kcal/mol)** | | | |
| Amber energy | -4895.571 | -5265.5 | -6490.194 |
| Restraints distance | 11.636 | 16.78 | 20.804 |
| Restraints dihedral angle | 2.229 | 2.14 | 4.606 |
| **RMSD* (Å)** | | | |
| Backbone (C’, N, Cα, O) | 0.271 | 0.324 | 0.268 |
| **RDC validation§** | | | |
| Total number of RDC values | 82 | 102 | 93 |
| RMS error | 0.948 (0.894) | 0.923 (0.916) | 0.904 (0.921) |
| Q-value | 0.319 (0.330) | 0.377 (0.397) | 0.397 (0.391) |
| **Ramachandran plot** | | | |
| Most favored regions (%) | 97 | 99 | 99 |
| Allowed regions (%) | 3 | 1 | 1 |
| Outliers (%) | 0 | 0 | 0 |

| Design ID | H6_fold-U _Nomur | H7_fold-K _Mussoc |
|---|---|---|
| PDB ID | 7BQS | 7BQR |
| BMRB Entry | 36339 | 36338 |
| **NMR distance and dihedral constraints** | | |
| **Detected NOE peaks** | | |
| $^{15}$N-edited NOESY | 1,727 | 2,184 |
| $^{13}$C-edited NOESY for aliphatic | 4,088 | 4,316 |
| $^{13}$C-edited NOESY for aromatic | 152 | 110 |
| **Distance constraints** | | |
| Total | 2,515 (100.0%) | 2,934 (100.0%) |
| Intra-residue | 436 (17.3%) | 484 (16.4%) |
| Inter-residue | | |
| Sequential (\|i-j\| = 1) | 436 (23.0%) | 663 (22.6%) |
| Medium-range (1 < \|i-j\| < 5) | 801 (31.8%) | 906 (20.9%) |
| Long-range (\|i-j\| ≥ 5) | 700 (27.8%) | 883 (30.1%) |
| **Dihedral constraints** | | |
| Total | 195 | 220 |
| **NMR structure statistics** | | |
| **Target function (Å$^2$)** | | |
| Total | 1.52±0.13 | 1.62±0.13 |
| Upl¶ | 0.0061±0.0007 | 0.0049±0.0007 |
| VDW | 6.8±0.40 | 7.4±0.4 |
| Dihedral | 0.60±0.05 | 0.52±0.05 |
| **Amber structure statistics (kcal/mol)** | | |
| Amber energy | -4861.6 | -5897.0 |
| Restraints distance | 28.792 | 18.584 |
| Restraints dihedral angle | 3.394 | 2.943 |
| **RMSD* (Å)** | | |
| Backbone (C’, N, Cα, O) | 0.174 | 0.185 |
| **RDC validation§** | | |
| Total number of RDC values | 141 | 95 |
| RMS error | 0.902 (0.776) | 0.913 (0.921) |
| Q-value | 0.405 (0.594) | 0.304 (0.391) |
| **Ramachandran plot** | | |
| Most favored regions (%) | 97 | 100 |
| Allowed regions (%) | 2 | 0 |
| Outliers (%) | 0 | 0 |

**Table S7. NMR and refinement statistics of 5 designed structures.**

¶ The Upl target function becomes more than zero when atom pairs are beyond the upper limits of distance constraints.

* The RMSD values were calculated for the 20 structures overlaid to the mean coordinates for the ordered regions, which were automatically identified by Fit_Robot using multi-dimensional non-linear scaling (*11*).
§ The RDC back-calculation was performed by PALES (*12*) using experimentally determined values of 1-bond $^{1}H$-$^{15}N$ RDC. The averaged correlation RMS and Q-value for NMR structure ensemble and design coordinates (as shown in parentheses) between the simulated and experimental values was obtained using the $^{1}H$-$^{15}N$ signals except completely overlapped peaks or the residues in low order-parameters (less than 0.8) predicted by TALOS+.

| | **H5_fold-0_Elsa** |
|---|---|
| **Data collection** | |
| Space group | $P2_1$ |
| Cell dimensions | |
| $a$, $b$, $c$ (Å) | 45.98, 33.66, 58.38 |
| $\alpha$, $\beta$, $\gamma$ (°) | 90.00, 93.11, 90.00 |
| Resolution (Å) | 45.9-2.33 (2.47-2.33)* |
| $R_{merge}$ | 0.090 (0.483) |
| $I/\sigma I$ | 13.63 (2.54) |
| Completeness (%) | 98.7 (92.3) |
| Redundancy | 6.4 (4.5) |
| **Refinement** | |
| Resolution (Å) | 45.9-2.33 |
| No. reflections | 49835 |
| $R_{work/}$ $R_{free}$ | 0.2066/0.2469 |
| R.m.s deviations | |
| Bond lengths (Å) | 0.003 |
| Bond angles (º) | 0.517 |
| Ramachandran plot statisties (%) | |
| Favored regions | 98.8 |
| Allowed regions | 1.2 |
| Outliers | 0.0 |

**Table S8. X-ray crystallography data collection and refinement statistics.**

*Statistics for the highest resolution shell are shown in parentheses. PDB ID: 7DNS.

| | RMSD between design and NMR (Å) | | $T_m$ (°C) |
|---|---|---|---|
| | **Cα atoms** | **Heavy atoms** | |
| **H5_fold-0_Chantal** | 1.9 | 3.8 | 118 |
| **H5_fold-0_Elsa** | domain-swapped | | 106 |
| **H6_fold-C_Rei** | 1.5 | 3.6 | 105 |
| **H6_fold-Z_Gogy** | 1.7 | 3.6 | 122 |
| **H6_fold-U_Nomur** | 3.1 | 4.4 | 116 |
| **H7_fold-K_Mussoc** | 2.2 | 4.0 | 139 |

**Table S9. Summary of experimental results for the six designs of the five folds.**

The second and third columns show the average RMSD between the design model and the 20 NMR structures using Cα atoms and heavy atoms respectively. The computationally designed regions were used for RMSD calculations. The last column shows the melting temperature $T_m$.

| | L1 | L2 | L3 | L4 | L5 | L6 |
|---|---|---|---|---|---|---|
| **H5_fold-0_Chantal** | BAABB/BAABB | GBB/GBB | E/E | GBB/GBB | - | - |
| **H5_fold-0_Elsa** | BAABB/BAABB | GBB/GBB | E/E | GBB/GBB | - | - |
| **H6_fold-C_Rei** | B/B | GBB/GBB | GB/GB | B/B | BAAB/BAAB | - |
| **H6_fold-Z_Gogy** | BGBB/BGBB | B/B | GBB/GBB | GB/GB | GBB/GBB | - |
| **H6_fold-U_Nomur** | BAB/BAB | BB/BB | AGB/AGB | AGBB/AGBB | AG/AGEB | - |
| **H7_fold-K_Mussoc** | A/A | AGABB/AGABB | AGB/AGB | AGBB/AGBB | BAB/BAB | AB/AB |

**Table S10. Comparison of ABEGO-based loop geometries of HLH motifs between design models and experimental structures for the six designs of the five folds.**

For the five designs except H5_fold-0_Elsa, the loop types for the design model (left) and the most frequent loop type in the NMR conformers (right) are shown. For H5_fold-0_Elsa, the loop types for the design model (left) and those of the crystal structure (right) are shown. The crystal structure of H5_fold-0_Elsa was domain-swapped due to the torsion angle difference of the residue immediately after the L4, Pro76, from A to B in the ABEGO representation.

**Table S11. Designed sequences and sequence identity among them.**

Computationally designed sequences are shown in uppercase and residues added to allow expression, purification, and the spacer between the designed sequence and the C-terminal His-tag are shown in lowercase. The rightmost column shows PSI-BLAST E-values against the nr database of nonredundant protein sequences.

#### H5_fold-0

| ID | sequence | E-value |
|---|---|---|
| Ana | mGEEEKRLKEILELIARWYEHVREKDRTGTEEDIVRKAVEEAAKTHGTNPKEVLERIARAIKKTGREQVARQAGTSEKTVEIIRRLWEREGslehhhhhh | 0.70 |
| Bill | mGEDEKELQRLLELLNRSYETEKRKDKSGSSKEWLRRAVEKAARQHGTSPSRLLELVQRAIEKKGRKELAKRMGTSEEDVKLLEELSKEQGslehhhhhh | 0.26 |
| Chantal | mGEEEKEIDKLVELFAQAYEDAREKKRNGTPEEWVRDAIEEAARRVGRSRSRVVEALRRYAEKHGKEELLKRAGITPEALKVIEKIEKEEGslehhhhhh | 0.73 |
| Chris | mGEEEKEIRKIVELFARAYKEAKEKKRNGTPEEWVRDAVEKAAQKVGRSRKDVVEALQKFADQEGEEELAKQLGISPEALKVIKKIRKEEGslehhhhhh | 0.65 |
| Danny | mGEEEKEIERLVELFAQAFREVKEKDQTGTPEEIARKAVERAAREEGRSRKRVVEALENYARKKGEEELLKRVGMTPEVWKVVQQIKKEEGslehhhhhh | 0.78 |
| Elsa | mGEEQKEIETLVELFAEAFREAKRQKKNGTPEEWARDAVEEAARQQGRSRKDVVEALTKYAQEQGRDELLKRLGITPEIYKVIQQIRKEEGslehhhhhh | 0.093 |
| Fay | mGEEQKELEKVLELFAQAFEQAKREKKNGTPEEWARDAVERAAQRVGRSRKDVVDLIEKAAREQGEKKLLERLGISPEVLKVIKKIQKEEGslehhhhhh | 0.46 |
| Flanklin | mGEEEKEREELIRLFAKAFEEYKRKHKNGSKEEIARRALERAARERGRSRSRVVEALEREARQTGEDELLKQVGMDPEVWKVVQQIKKEEGslehhhhhh | 1.6 |
| Hermine | mGDEDEKKKRILEALLRAYEEWRKKDKTGSQEDVVRKAVERAAREAGTNPRDVLRVVQEEIEKTGPDELAKKIGTSPSVVELLRKVYEEEGslehhhhhh | 0.91 |
| Larry | mGEREKKIKEVVRRFAKAYEEVRKTNQNGSEKEIVERAVREVAKEEGTSPREVLKILVEFIKRKGPEEFAKEAGSSSEAAKIIEELLRREGswslehhhhhh | 2.0 |

| | Ana | Bill | Chantal | Chris | Danny | Elsa | Fay | Franklin | Hermine | Larry |
|---|---|---|---|---|---|---|---|---|---|---|
| Ana | 1 | 0.444 | 0.4 | 0.411 | 0.378 | 0.3 | 0.333 | 0.311 | 0.489 | 0.444 |
| Bill | 0.633 | 1 | 0.444 | 0.411 | 0.422 | 0.4 | 0.4 | 0.422 | 0.467 | 0.367 |
| Chantal | 0.578 | 0.656 | 1 | 0.722 | 0.656 | 0.667 | 0.611 | 0.567 | 0.344 | 0.4 |
| Chris | 0.589 | 0.611 | 0.822 | 1 | 0.611 | 0.678 | 0.667 | 0.544 | 0.378 | 0.4 |
| Danny | 0.556 | 0.6 | 0.756 | 0.733 | 1 | 0.689 | 0.589 | 0.733 | 0.378 | 0.378 |
| Elsa | 0.5 | 0.556 | 0.767 | 0.756 | 0.767 | 1 | 0.689 | 0.611 | 0.311 | 0.322 |
| Fay | 0.511 | 0.578 | 0.744 | 0.767 | 0.7 | 0.767 | 1 | 0.556 | 0.322 | 0.278 |
| Franklin | 0.544 | 0.578 | 0.7 | 0.711 | 0.789 | 0.733 | 0.678 | 1 | 0.367 | 0.367 |
| Hermine | 0.7 | 0.644 | 0.556 | 0.611 | 0.589 | 0.511 | 0.533 | 0.589 | 1 | 0.367 |
| Larry | 0.589 | 0.544 | 0.544 | 0.522 | 0.556 | 0.489 | 0.444 | 0.522 | 0.578 | 1 |

### H6_fold-C

| ID | sequence | E-value |
|---|---|---|
| Cyrus | mGSSVERAAEEVFQIIQQSPEVFERLIESMKEWMKKQGSSPDELRKLEKDLKEARKRAEEQKRQGNEEEMRRIVQKVLKSPAFKQAVQLMEEQEPNNPEVKKLKEAMEEVENGslehhhhhh | 0.28 |
| Dan | mGDEAKKAAEEMIKELSKNPELMKRLIKEMEKWLKQKGVSPDELERTRQEMETRIKEAIERKKRGDEESAREVMEEVLKSPAFKQAARNLEEQEPNNPEAQKMRELAERAEKGslehhhhhh | 418 |
| Max | mGEEAKKAAEEVLQLMDKSPDLMQQVIEEMKKVWKNKGASPEQLEKQERKVKTLVEEARKRKEEGNEEDARKVMLEVLKSPAFKQAVKRVEEQEPNNPEAKRLLELAERASQGslehhhhhh | 91 |
| Melvin | mGDDAKKMARKVLEIVTKSPDVMKTLIKRAEERLEKEGRSDEDKQRWRETIKKKVQKAIELQKEGNEEKAEETMEEVLKSPAFKQAVEMLEEQEPNNPEAQTLRRAMEEAERGslehhhhhh | 1.8 |
| Moca | mGDDAKKRARDLARTIAESPEVAERLAQALREKLENQGQSEDEIRSRVTELRKRLEEAIRQWRTGNKDEAMRVALEVLKSPAFKQAVEALEEQEPNNPEARKLLQMAEDVEKGslehhhhhh | 0.047 |
| Paul | mGQEARDAAREMLKEVERSPSVMKTVVEMARERAREEGQSPDQLKEIAREIEENVRRALELWRRGNSEKAEKVMEKVLKSPAFKQAMEMLEEQEPNNPEMNKVIELAKRASRGslehhhhhh | 0.27 |
| Rei | mGDEAEKQAERALELVRKSPDLLKKLLEAMAEELKRQGKSPDEIQKAKDEVKTKVEQAIREWKQGNEEQARKDMRKVLKSPAFKQAVKVMEEQEPNNPEVQELKKAMEEAERGslehhhhhh | 2.4 |

| | Cyrus | Dan | Max | Melvin | Moca | Paul | Rei |
|---|---|---|---|---|---|---|---|
| Cyrus | 1 | 0.455 | 0.473 | 0.42 | 0.455 | 0.402 | 0.509 |
| Dan | 0.58 | 1 | 0.571 | 0.518 | 0.455 | 0.491 | 0.491 |
| Max | 0.616 | 0.705 | 1 | 0.464 | 0.402 | 0.473 | 0.527 |
| Melvin | 0.545 | 0.643 | 0.545 | 1 | 0.455 | 0.5 | 0.545 |
| Moca | 0.58 | 0.589 | 0.607 | 0.589 | 1 | 0.402 | 0.446 |
| Paul | 0.58 | 0.634 | 0.616 | 0.616 | 0.58 | 1 | 0.429 |
| Rei | 0.652 | 0.643 | 0.607 | 0.705 | 0.589 | 0.571 | 1 |

### H6_fold-Z

| ID | sequence | E-value |
|---|---|---|
| Gogy | mGDERKLEEVTEEMRKMAENMDGQDPEKVKEIVRRALQQMANDNPEVSEQLRELAKRKGTSPSEVIKDLAEQVWRAMERAREGDKDTARELIRKFADDLGISPEQVKKFIKIMREVQRKEDGslehhhhhh | 0.065 |
| Hoto | mGDDDEIRRILELLEKIARSLDGASEDTIREVIEQAIRDLQKQNPEFKKQVERVAKEQGTSPDEVIKKIVEIIWEAAERVRRGNKDEAEKVIRKLEKELGISPSTVRQLMEEIVRVLKEKKGslehhhhhh | 0.0003 |
| Kome | mGTDEELKKVDEELRRIQTQLDGQTEDEIRRIMEKVMKQLTKTNPEFKKQVKRVAEEQGTSPSEVLKEIIERLEKAAQEAKKGNTEEAQKIVKELAKTLGVTWSEIRRILEQIREEIERTQGslehhhhhh | 0.35 |
| Nazu | mGDDQELEEIMRLAEDVSREVKGQSPDELKKMMKKLLEELTKENPEFKETIRENAEKEGTSPSEVWRKVIERVEQIMRDVTKGDKEQARKEIEELRKELGVSLKDIKELMKRLKEVVKKSRGslehhhhhh | 0.002 |
| Seri | mGESDEIKKILQELEKIWQNIKGVSDEQIQEVFKQLVKELTREDPQFEKSVRRLAKEEGTSPSNVQEKLLKLIKDAAENVRRGNKDEAEKNIREIAKNLGVSHSDVLRIVKTLRELLEQAKGslehhhhhh | 0.015 |
| Siro | mGSSEDARTVVELLQKIRQEIEGLSPDQIKEVVKRAIDELEKENPQVSKNLEEKAQKQGTSPSKLKEKVAELIREAAERAQNGDSEEAKKLVEKIARILGISWKDVVRMMKEIIRRLRDENGslehhhhhh | 0.018 |
| Suzu | mGSKETLKKILKTLQEAKKELKGTSPEEIKKRIVKLLKEVAKESPEIKKNIKEEAERQGTSPSEVLKEIADLIREAMSRAEEGNSEEAEKMIDEIARIMGISWDQVLEILDRLVEQLKKDRGslehhhhhh | 0.054 |

| | Gogy | Hoto | Kome | Nazu | Seri | Siro | Suzu |
|---|---|---|---|---|---|---|---|
| Gogy | 1 | 0.364 | 0.289 | 0.347 | 0.306 | 0.364 | 0.322 |
| Hoto | 0.554 | 1 | 0.446 | 0.364 | 0.471 | 0.421 | 0.355 |
| Kome | 0.57 | 0.661 | 1 | 0.372 | 0.372 | 0.314 | 0.397 |
| Nazu | 0.545 | 0.554 | 0.595 | 1 | 0.347 | 0.331 | 0.397 |
| Seri | 0.463 | 0.645 | 0.62 | 0.554 | 1 | 0.397 | 0.38 |
| Siro | 0.545 | 0.612 | 0.554 | 0.529 | 0.579 | 1 | 0.413 |
| Suzu | 0.529 | 0.57 | 0.62 | 0.595 | 0.521 | 0.636 | 1 |

### H6_fold-U

| ID | sequence | E-value |
|---|---|---|
| Cacyd | mGETDIRAALTALKTIIKKASNPEVAEAAKRLIEALKKASPEQRERAARKLIKALKALAKGDTERAQKLIKEAAEEAGLSPEDIKKLQKAARWLKEQGLVREALQAAQDVQDKGslehhhhhh | 0.05 |
| Hatake | mGETDIRAAQEALKVIIKKASNPEVAEAAKSLLEALKKASPEQRRRAAEKIIKALKALAKGDTERAVELLREAAEEAGLSPREIEKLRKAARWLKEQGLVREALRRAQEVQDNGslehhhhhh | 0.72 |
| Kazi | mGETERRAAERALEVLIRRATDPELQQAVKDLRKVLNTASPERQKRAAEKIFRALEAAAQGNKEKAKKLAEEAARVAGASPEIIRRLQKAVEKAQKRGIVKEARKAAQEVKENGswslehhhhhh | 0.041 |
| Morit | mGDREIKAALTALKVIIKKASNPEVAEAAERLLRALKKASPEQRRRAAENIIRALKALAKGDTERAAKLLKKAAEEAGLSPDTIKKLKKAARWLKEQGLVREALKSAKEVTKKGslehhhhhh | 0.004 |
| Nomur | mGETKAKAAQEALRAAREQATTPEAQKALEELEKVLKTASPEQWRQAAEKIFEAFREASNGNTEKAKKLLEEAARTAGASPEIIKKLASALERLAEEGAAKEAARQAEEVRKRGslehhhhhh | 0.35 |
| Sait | mGDKEAQAAQEAIKTAIKSATNPEAQRALKEFSRVANEASPDQWRKAADLIRKALEEASRGNTERAERLARKAAKVVGASPEIIERLARAIRELARTGAAKKARQVAEKVKKKGslehhhhhh | 0.051 |
| Sho | mGEEVLKAAKRALEVIIRRATDPELQQAVKELQKILSTASDERRERAAKEIYRAAEAAAQGNKEEARKRLEKAAKELGASPEQIEKAKKALDEAQKRGIVKEAKEEAEKQQRKGswslehhhhhh | 0.15 |
| Takin | mGDTEAKAASRALQTIIDQATDPELKKALEDLRRRAEEASPDQWRQIAKAILKALELLDRGNTEEARKELEKAARKAGASPEVAKKLQKAFSEATKQGIAKEASKEAQKVRDKGslehhhhhh | 0.47 |

| | Cacyd | Hatake | Kazi | Morit | Nomur | Sait | Sho | Takin |
|---|---|---|---|---|---|---|---|---|
| Cacyd | 1 | 0.823 | 0.487 | 0.779 | 0.442 | 0.416 | 0.434 | 0.451 |
| Hatake | 0.876 | 1 | 0.496 | 0.779 | 0.487 | 0.434 | 0.434 | 0.425 |
| Kazi | 0.619 | 0.619 | 1 | 0.46 | 0.549 | 0.46 | 0.628 | 0.513 |
| Morit | 0.858 | 0.841 | 0.593 | 1 | 0.469 | 0.434 | 0.451 | 0.478 |
| Nomu | 0.531 | 0.54 | 0.646 | 0.558 | 1 | 0.54 | 0.451 | 0.513 |
| Sait | 0.531 | 0.531 | 0.611 | 0.558 | 0.664 | 1 | 0.407 | 0.504 |
| Sho | 0.54 | 0.522 | 0.726 | 0.549 | 0.531 | 0.558 | 1 | 0.504 |
| Takin | 0.566 | 0.522 | 0.637 | 0.558 | 0.593 | 0.593 | 0.566 | 1 |

### H7_fold-K

| ID | sequence | E-value |
|---|---|---|
| Chario | mGDEEREKLRKIARKALKDAASEAKKRGIITPEAIERIANLLADAAQEWKEGNEERAEKL LRKAAKVFVEAAKKTGASADEARRVLERIRKALSNAPSDELRAIAKSPRLKKAIKEAIEEI RRIEEETGslehhhhhh | 0.49 |
| Dark | mGDDERKKIKEAAREAVERALEDAKRQGIVSPESLERIAEKIARAAEALEKGNSEKARKLI EEAARTILKEAERAGASSRWVEEILERLRQALEKAPSPRLQKLAQSPVFKTALEKIIREAR KREKKRGslehhhhhh | 0.002 |
| Jazzbird | mGDDDRKKLEQTVEEAIRKALEDAVRSGAITPEAAERIAQEIAEAAAQWRQGNTEEAEK AARRAIKVLLEAARKSGASSDQISELLERIREALSNAPSPEVQQLAKSPRFQQVLKEAKKE AKRIEKETGswslehhhhhh | 0.000002 |
| Mussoc | mGDEDKEKLKREAERALSEALSEFEKQGKITPETLKRLAEEIAEAALAQQQGDSERLEKA ARRFAETLLRALKESGASAEEIEEAIERIRKALSKAPSPQLQKLANSPQWQTALQEAIKKA RQEKKEKGslehhhhhh | 0.64 |
| NewWave | mGGSDRSDELRKQAEEAARKAFEEAKKRGVITPDAIKKVAEEIARAAELWRQGNTEEAE KAVRNAIKVILEAAKKSGASSQEVSDALERLRQALENAPSPEVQQLAKSPRFQQVIEEAEK EAKRIEKETGslehhhhhh | 0.043 |
| R3 | mGDDDREKLRETAREALREALSKFEKQGKITPEVLERLAEAIAEAASALEKGDSERLEKA ARKFAQILLESLKKSGASAEEIEEALRLAQKALSKAPSPQLQRVANSPRWQEALNEAIKRA REEEKRTGslehhhhhh | 0.32 |
| Rush | mGEDEEKKIEEAARQAVETALEDARRQGQITPERIEKIAEKIARAATALRRGNKEEAIKLI KEAAQIIVEAARESGASSEWIEEVLRRIEEALRRAPSPQLQKLADSPAFQKAAREAIERAR EEEERTGswslehhhhhh | 0.079 |
| Third | mGDDEAKKIEDAAREAVKKALSEALKRGTITPDVIEEIAQRLAEAAIALEEGNKEKAAKL AKEALAKLIEEAAREGASPRWIERLIDRIEKQLANAPSPQLQKLAESPVFKKVLEEAKQRA DEVRRKTGslehhhhhh | 0.028 |

| | Chario | Dark | Jazzbird | Mussoc | NewWave | R3 | Rush | Third |
|---|---|---|---|---|---|---|---|---|
| Chario | 1 | 0.422 | 0.531 | 0.461 | 0.531 | 0.484 | 0.461 | 0.453 |
| Dark | 0.578 | 1 | 0.469 | 0.453 | 0.453 | 0.477 | 0.594 | 0.516 |
| Jazzbird | 0.68 | 0.602 | 1 | 0.508 | 0.688 | 0.516 | 0.516 | 0.477 |
| Mussoc | 0.594 | 0.617 | 0.609 | 1 | 0.453 | 0.711 | 0.453 | 0.406 |
| NewWave | 0.656 | 0.625 | 0.773 | 0.594 | 1 | 0.445 | 0.453 | 0.406 |
| R3 | 0.594 | 0.617 | 0.609 | 0.781 | 0.562 | 1 | 0.523 | 0.445 |
| Rush | 0.602 | 0.695 | 0.625 | 0.594 | 0.578 | 0.664 | 1 | 0.555 |
| Third | 0.578 | 0.656 | 0.617 | 0.539 | 0.539 | 0.594 | 0.656 | 1 |
